# Fecal extracellular vesicles from adults with autism spectrum disorder affect mouse behavior and gut motility by targeting the neuronal nitrergic pathway

**DOI:** 10.64898/2026.09.28.754921

**Authors:** Baptiste Ganachaud, Irem Buruk, Catherine Le Berre-Scoul, Philippe Aubert, Camille Trottier, Néfis Djelassi, Mathéus Moreau, Florian Petrilli, Jean-Romain Richard, Camille Dupuy, Antoine Lefevre, Patrick Émond, Lydie Nadal-Desbarats, Marine Gilles, Benoît Quinquis, Florence Thirion, Joel Doré, Ramaroson Andriantsitohaina, Marion Leboyer, Sophie Talon, Angela Tesse, Justine Marchix, Michel Neunlist, Martial Caillaud, Hélène Boudin

## Abstract

Beside behavioral symptoms, many patients with autism spectrum disorder (ASD) present gastrointestinal disorders and gut microbiota dysbiosis. In order to better understand the mechanisms underlying microbiota-gut-brain interactions in ASD, we isolated fecal extracellular vesicles (EVs), as a source of microbiota-derived EVs, from control individuals (f-EV-CTL) and patients with ASD (f-EV-ASD) to study in mice their impact on brain and gut. The f-EV-ASD induced hypersocial and anxiolytic-like behavior, and reduced gastrointestinal transit time compared to mice treated with f-EV-CTL. In the colon, f-EV-ASD altered motility through the enteric neuronal nitric oxide (NO)-sensitive component. f-EV-ASD potentiated NO signaling and disrupted intracellular calcium dynamics in enteric neurons. This effect was mediated by butyrate and valerate through FFAR3 receptor. Our findings establish fecal EVs as critical mediators of microbiota-host communication, linking microbial metabolites to altered behavior and gut function and identifying a potential mechanism underlying ASD pathophysiology.

## INTRODUCTION

Autism spectrum disorder (ASD) is a heterogeneous neurodevelopmental condition characterized by deficits in social interactions, repetitive behaviors, and restricted interests^1^, which currently affects 1.2% of the global population^2^. Beyond behavioral symptoms, gastrointestinal (GI) symptoms are among the most frequently associated comorbidities in ASD (McElhanon et al., 2014). Nearly 50 to 80% of individuals with ASD experience GI disorders, including diarrhea and constipation^3,4^. Despite the high prevalence of GI disorders and their impact on ASD severity, the pathophysiological mechanisms underlying GI dysfunction in ASD remain poorly understood.

A central player in gut physiology is the enteric nervous system (ENS), which is distributed throughout the gastrointestinal tract and controls epithelial barrier permeability and gut motility^5^. Alterations in neuronal connectivity or in the proportion of cholinergic and nitrergic neurons within the ENS are responsible for transit disorders^6–8^. Mouse models of ASD also exhibit GI disorders, associated with ENS alterations characterized notably by changes in the proportion of nitrergic neurons^9,10^. However, the mechanisms involved in GI disorders and ENS modifications remain unknown but may involve the gut microbiota.

Among the factors potentially involved in ASD pathophysiology, the gut microbiota has emerged as key contributor. Changes in the composition of the gut microbiota have been described in children and adults with ASD^11–15^. Moreover, the transfer of gut microbiota or fecal supernatant from individuals with ASD to mice induces ASD-relevant behavioral disorders, GI disorders, and remodeling of the ENS, thereby suggesting a role for the gut microbiota in both central and intestinal symptoms^11,16,17^. However, the mechanisms and bacterial mediators involved in the effects of the microbiota on host organs, especially the gut and the brain, remain to be defined in ASD.

Bacteria-derived mediators include bacterial metabolites, components of the bacterial cell envelope and cell wall, and extracellular vesicles (EVs). EVs are of particular interest because they can encapsulate and transport diverse signaling molecules, including proteins, lipids, and nucleic acids, thereby facilitating communication with recipient cells and organs both locally and at distant sites^18–20^. EVs isolated from probiotic or commensal gut bacteria have been shown to modulate gastrointestinal and brain functions following administration to mice^21–23^ and to reverse the molecular and behavioral alterations in a mouse model exhibiting ASD-like phenotypes^24^. The contribution of EVs originating from complex microbial communities, such as the gut microbiota, has also been investigated in the context of liver disease, using fecal samples as a source of microbiota-derived EVs. Fecal EVs (f-EVs) isolated from patients with liver disease reproduced several disease-relevant pathological features when transferred to mice^25,26^. However, whether f-EVs contribute to the behavioral and gastrointestinal dysfunctions associated with ASD remains to be determined.

To address this question, f-EVs were prepared from stool samples, which predominantly contain bacteria-derived EVs^27^. We identify f-EVs as previously unrecognized mediators of microbiota-host communication in ASD. Transfer of ASD-derived f-EVs into mice altered social and anxiety-like behaviors, modified synaptic protein expression in the brain, and impaired gut motility through nitrergic neuronal signaling. Enteric neurons emerged as direct f-EV targets, with their Ca²⁺ dynamics controlled through a butyrate–FFAR3-dependent mechanism. These results establish a mechanistic link among microbial EV signaling, enteric neuronal activity, and ASD-associated brain–gut dysfunction.

## MATERIAL AND METHODS

### Subject recruitment and sample collection

Individuals with ASD (n=30) were recruited through the Créteil Autism Expert Center (Chenevier–Mondor Hospital in Créteil, France). The participants, all above 18 years, were diagnosed with ASD without intellectual disability according to the fifth edition of Diagnostic and Statistical Manual of Mental Disorders (DSM-V) and the Autism Diagnostic Observation Schedule -2 (ADOS-2) completed with the Autism Diagnostic Interview-Revised (ADI-R) (Table 1). Gastrointestinal symptoms were assessed through questionnaire incuding 7 items from Rome IV questionnaire including the Bristol Stool Scale. Individuals who had received antibiotics or antifungal agents during the previous two months were excluded from the study. Neurotypical controls (n=15) were recruited from the Center for Clinical Investigation (Henri Mondor Hospital, Créteil, France). The inclusion criteria were an age > 18 years and no psychiatric or somatic medical history. The non-inclusion criteria were: smoking ≥ 5 cigarettes per day; taking psychotropic medication; BMI > 24.9 and gastrointestinal disorders (Table 2). The participants were enrolled in the study after signing an informed consent form. This study was approved by the French Ethical Committee for Biomedical Research (Comité de Protection des Personnes, Ile de France 2, protocole 2017-A01318-45 and used a collection registered under no. 16113 with the collection code DC 2009-930, approved by CCP Ile de France V). The fecal samples were collected at home by each participant, stored at 4-8°C under anaerobic atmosphere using the GENboxanaer kit (Biomérieux, Marcy L’Etoile, France), and carried to the Centre d’Investigation Clinique of Créteil Hospital in less than 24h. Fecal samples were then aliquoted and stored at -80°C.

**Table 1:** Summary of demographic characteristics and gastrointestinal symptoms of CTL individuals and ASD patients. Age comparison CTL vs. ASD, p=0.55, Student ’s t-test.

|  | CTL | ASD |
| --- | --- | --- |
| **Number of subjects** | 15 | 30 |
| **Minimum-Maximum age**<br>(Mean age ; 1st-3rd quartile) | 20-49<br>(32 ; 24-39) | 18-45<br>(31 ; 24-36) |
| **Gender** |  |  |
| Number of Female | 6 (40%) | 11 (37%) |
| Number of Male | 9 (60%) | 19 (63%) |
| **Gastrointestinal symptoms** |  |  |
| **Low** | NA | 12 (40%) |
| **Medium/High** |  | 18 (60%) |

### Microbiota analyses

DNA extraction from aliquots of fecal samples from 26 individuals with ASD and 10 neurotypical controls was performed following the MetaGenoPolis procedure SOP001 v1 (https://mgps.eu/sops/mgp-sop-001-v1/) based on IHMS recommendations. DNA was quantified using Qubit Fluorometric Quantitation (ThermoFisher Scientific, Waltham, US) and qualified using DNA size profiling on a Fragment Analyzer (Agilent Technologies, Santa Clara, US). Three micrograms of high molecular weight DNA (>10 kbp) were used to build the library. Shearing DNA into fragments of approximately 150 bp was performed using an ultrasonicator (Covaris, Woburn, US) and DNA fragment library construction was performed using the Ion Plus Fragment Library and Ion Xpress Barcode Adapters Kits (ThermoFisher Scientific, Waltham, US). Purified and amplified DNA fragment libraries were sequenced using the Ion Proton Sequencer (ThermoFisher Scientific, Waltham, US), generating a minimum of 20 million reads of 150 bp per sample. Raw reads were quality trimmed using AlienTrimmer v2.0 with the following parameters: “-k 10 -l 45 -m 5 -p 40 -q 20”. Human-related reads were removed using Bowtie2 v2.5.1 and the Human genome reference genome T2T-CHM13v2.0 (accession GCF_009914755.1) and a nucleotide identity threshold of 90%. Samples were taxonomically profiled with Meteor2^28^ (v2.0.18, normalization = “fpkm”, rarefying = 20 million reads), using both the human gut and human oral catalogues available through Meteor2. The Metagenomic Species (MSP) abundance tables generated using each catalogue were subsequently merged into a single MSP abundance table.

### Preparation and characterization of f-EVs

The preparation of f-EVs was carried out as previously described^25^. Feces were thawed on ice and incubated overnight in phosphate bufferred saline (PBS, BIOSERA, Cholet, France). The fecal suspension was then sequentially centrifuged at 700 g for 15 min, 3600 g for 15 min, and 10,000 g for 30 min with the supernantant collected after each centrifugation step. The last supernatant was then filtered through a 0.22 µm filter and centrifuged at 17,000 g during 1h. The resulting pellet was resuspended in PBS and ultracentrifuged at 150,000 g for 2h at 4°C. The pellet was then resuspended in PBS overnight and ultracentrifuged again at 150,000 g for 2h at 4°C. The pellet was resuspended in 200 µL of sterile PBS and stored at -80°C until use. Protein concentration was determined using the BCA protein assay (Sigma-Aldrich). The size and number of particles was characterized by Nanoparticle tracking analysis (NanoSight NS300, Malverne Instruments, Orsay, France). Transmitted electron microscopy of f-EV preparations was performed with a JEM-1400 microscope (JEOL, Tokyo, Japan) at a voltage of 120KV. The images were acquired with a Orius 832 camera from Gatan Inc. (Pleasanton, CA 94588, USA).

### Metabolomics analyses of f-EV preparations

#### Targeted metabolomics

For sample preparation, 800 µL of methanol was added to 25 µL of EV’s extract. The samples were vortexed and centrifuged at 15,000g for 15 min at 4°C.

Following centrifugation, 750 µL of the resulting supernatant was carefully collected and transferred to a 96-well plate. For C18 and HILIC analyses, 350-µL aliquots were separately evaporated to dryness under a nitrogen stream at 40°C for 30 min and subsequently reconstituted in 100 µL of acetonitrile/water (2:8, v/v) for C18 analysis or acetonitrile/water (8:2, v/v) for HILIC analysis. A pooled quality-control (QC) sample was prepared by combining 10 µL of each sample extract. LC-HRMS analyses were performed according to the parameters previously reported [1]. C18-based chromatography was performed in both positive and negative ionization modes, while HILIC chromatography was performed only in positive ionization mode. Data processing and feature annotation were performed according to the methodology previously described^29^.

#### Short chain fatty acids (SCFA)

A volume of 10 µL of EVs was added to a 96-Deepwell plate. Then, 40 µL of methanol was added to each well, and the plate was sealed with an aluminum cover. The plate was shaken on a planar shaker at force 7 for 5 minutes. Afterward, 3-nitrophenylhydrazine (3-NPH) derivatization was performed according to previously published method^30^. The injection volume was set to 15 µL into the LC-MS/MS system.

#### LPS

LPS levels were determined using the Pierce Chromogenic Endotoxin Quantification Kit (ThermoFisher, #A39552), following the manufacturer’s instructions.

### Animals

All protocols were performed according to the the Animal care and use committee of Nantes (France) and were approved by the French Ministry of Research (Agreement Apafis NO. 47154-2024012614323169 v5). C57BL/6Rj mice (7 weeks old) were obtained from Janvier Laboratory (Le Genest Saint-Isle, France) and were housed in 12h light-dark cycle with free access to water and food. After 10 days of habituation, mice were gavaged with 5.10^8^ f-EVs/g of mouse weight diluted in 100 μl of PBS every two days for two weeks (Fig. 2A). Each mouse corresponded to a single individual.

### Behavioral tests

#### Three-chamber social preference test

Mice were tested in a three-chamber apparatus consisting in a central compartment connected with two lateral compartments. First, the test mouse was acclimated to the empty apparatus and allowed to freely explore for 10 min. Then, an object was placed in a lateral compartment (nonsocial stimulus) and an unfamiliar mouse, matched for age and sex, was positioned in the other lateral compartment (social stimulus). The tested mouse was then placed in the central compartement and allowed to explore the entire arena for 10 min. The time spent in each compartment was determined by videotracking using ViewPoint Software (Lyon, France).

#### Marble burying test

Mice were individually placed in large cages (30 × 20 cm) filled with 5cm of wood chip bedding during 10 min for habituation. Mice were temporarily removed from their cages, and 20 glass marbles were evenly distributed on the surface of the bedding. The mice were then returned to their respective cages and allowed to explore for 10 min. The number of buried marbles was independently counted by three experimenters blinded to the experimental conditions. A marble was considered buried when at least two-thirds of its volume was covered with bedding.

*Elevated plus maze.* The apparatus consisted of two open arms and two closed arms (30 × 8 x15 cm) connected by a central platform (8 × 8 cm) and elevated 50 cm above the floor. Mice were placed in the center of the maze facing an open arm and allowed to explore for 5 min. Time spent in each arm was automatically quantified using videotracking. The percentages of the numbers of entries in open arms were calculated as entries in open arms × 100/(entries in open arms + entries in closed arms).

### Intestinal barrier permeability

#### In vivo

Mice received an oral gavage of a 100 µL solution containing 10 mg/mL of Fluorescein–5.6 sulfonic acid (FSA; F1130, Invitrogen, Life Technologies, Carlsbad, CA, USA), 10 mg/mL of TRITC-dextran (TD4, TdB Labs, Uppsala, Sweden), and 10 mg/mL of horseradish peroxidase (HRP; P8250, Sigma-Aldrich), diluted in 0.5% carboxymethyl cellulose. Blood samples were obtained from the tail vein 60 min post-gavage and the plasmatic concentration of FSA and dextran was determined by measuring the fluorescence intensity using a microplate spectrofluorometer (Varioskan). The quantification of HRP was conducted by measuring its enzymatic activity using 3,3′,5,5′ tetramethylbenzidine (TMB, BD Biosciences) as substrate.

#### Ex vivo

Mice were euthanized by cervical dislocation and segments of ileum and colon were dissected and mounted in Ussing chambers (Transcellab, TBC, Paris, France) in 2 ml of Ham’s Nutrient Mixture (Life Technologies) continuously oxygenated (95% O2 and 5% CO2) and maintained at 37°C. After 30 min of equilibration, a mixture of FSA (1 mg/ml), TRITC-Dextran (10 mg/ml) and HRP (3.75 mg/ml) was added at the apical side. The fluorescence levels of 150 μl aliquots taken from the basolateral side were measured at 30 min intervals over 180 min using a spectrofluorometer. To assess permeability to HRP, enzymatic assay was performed using TMB substrate. Permeability was determined by calculating the slope of fluorescence intensity over time for FSA and TRITC-dextran and of HRP activity measured over time in the specimens, using a linear regression fit model measured in the specimens (GraphPad Prism Software).

### Gut motility

#### In vivo

To assess fecal pellet output (FPO), mice were placed individually in a clean cage without bedding, food and water. Fecal pellets were collected during 2h and counted every 15min. The % of water content of the feces was assessed as the difference in weight between dried and fresh feces and divided by the weight of fresh feces. To measure the total transit time, mice were gavaged with 100 µl of a carmine red solution (60 mg/ml) suspended in a 0.5% carboxymethylcellulose solution. Total transit time was defined as the interval between the initiation of gavage and the time of first detection of carmine red in feces.

#### Ex vivo

Segments of ileum and distal colon were placed in an organ chamber (Radnoti, California, USA) with 15 mL of Krebs solution (NaH_2_PO_4_.2H_2_O 0.187 g/L, NaCl 6.84 g/L, KCl 0.35 g/L, NaHCO_3_ 2.10 g/L, glucose 1.98 g/L, CaCl_2_.2H_2_O 0.368 g/L, MgCl2.6H_2_O 0.244 g/L, pH 7.4) at 37°C and continuously bubbled with 95% O_2_ and 5% CO_2_. The contractile responses were recorded using isometric force transducers connected to an AD Instruments PowerLab 8/30 data acquisition system and the Labchart data analysis software. Tissues were stretched to an initial tension of 1 g, and after a 60-minute equilibration period, were subjected to electrical field stimulation (EFS) employing a STG 4008 MCS electrical stimulator (Multi Channel Systems, Reutlingen, Germany). The parameters established for the EFS included a train duration of 10 s, a pulse frequency of 20 Hz, a pulse duration of 400 µs, and a pulse amplitude of 11 V. This methodological procedure was replicated 3 times, with a 4-minute interval between each repetition. Subsequently, nitro-L-arginine methyl ester (L-NAME, 50 µM; N5751, Sigma-Aldrich), which acts as an inhibitor of nitric oxide synthases (NOS), was introduced into the baths following an incubation period of 10 min, after which atropine (1 µM; A0257; Sigma-Aldrich, Merck, Darmstadt, Germany), a muscarinic receptor antagonist, was administered. The identical EFS stimulation protocol was used. At the end of the experimental procedure, tissues were exposed to sodium nitroprusside (SNP, 71778-25G, Sigma-Aldrich, 100 µM), a nitric oxide (NO) donor, and tissue contractions were recorded 1 min subsequent to SNP addition.

### NO measurements

NO concentration was evaluated by electron paramagnetic resonance using Fe^2+^ diethyldithiocarbamate (Fe(DETC)2) as spin trap. A solution of Na-DETC, 15 mM in Kreps-HEPES, was mixed to a solution of FeSO4-7H2O, 8 mM in water, and was immediately used to incubate mouse pieces of organs or enteric neuron cultures. After 45 min incubation at 37 °C, spin trap was removed, and the samples were frozen in liquid nitrogen and analyzed in a Dewar flask at 77°K using an EPR Miniscope MS5000 (Bruker, Germany). The signals were quantified by measuring the peak amplitude from the obtained spectra, and were expressed in arbitrary units (AU). Data were normalized to dry weight for colon samples and to the total amount of proteins for cerebral tissues and cultured enteric neurons.

### Western blot

Samples were lysed in RIPA buffer containing phosphatase and protease inhibitors. After centrifugation at 10,000g for 10min at 4°C, samples (10-30 μg) were loaded on a NuPAGE 4%– 12% Bis-Tris Acrylamide gel with MES SDS running buffer. Proteins were transfered to nitrocellulose membranes using the iBlotTM Dry Blotting System (7 min at 20 V). Membranes were then blocked in TBS-Tween 5% milk and incubated overnight at 4°C with the following primary antibodies: rabbit monoclonal anti-beta-3-tubulin (1:1,000; Cat: ab52623, Abcam), mouse monoclonal anti-PGP 9.5 (1:500, MA1-83428, Thermo Fisher Scientific), mouse monoclonal anti-PSD95 (1:1000, MA1-045, ThermoFisher), mouse anti-nNOS (1:1000, 610308, BD Biosciences), rabbit anti-phospho-nNOS (Ser1417) (1:1000, PA1-032, ThermoFisher Scientific), mouse anti-3Nitrotyrosine (1:500, ab110282, Abcam), anti-β-actin antibody (1:5,000; A5441, Sigma). Membranes were then washed and incubated for 1 h with the appropriate HRP-conjugated secondary antibody diluted in TBS containing 5% milk. Proteins were detected by chemiluminescence with Clarity Western reagents using a ChemiDoc XRS+ system Bio-Rad. The intensity of signal was analyzed using ImageJ software (https://imagej.net). Data are expressed as relative values to β-Actin signal.

### Enteric neuron cultures and treatment

Cultures of rat enteric neurons were performed as previously described^31^. Briefly, embryos were collected from pregnant Sprague-Dawley rats on gestational day 15 (Janvier Labs, Le Genest Saint-Isle, France) and the whole intestine was dissected, cut into 8 equal-length pieces and placed in a 24-well plate precoated with collagen (0.1 mg/mL). After 5 days of culture in enteric glia-conditioned neurobasal medium supplemented with B27, 10 mM Glutamine, and GDNF (50 ng/mL), cells were collected with 300 µl of accutase. Cell suspension was then centrifuged (1,500 rpm, 5 min) and the pellet was resuspended in the enteric glia-conditioned medium + GDNF (50 ng/ml). Cells were plated in 24-well plate for NO measurements, or in a 48-well plate for western blot, or in Ibidi 8-well plate for Ca^2+^ imaging and immunofluorescence. The next day (Day 1), the antimitotic agent cytosine arabinoside (5 µM) was added to prevent proliferation of glial and muscle cells. Enteric neuron cultures were treated on Day 11 and 12 with f-EVs (10^9^ particles/ml) and analyzed on Day 13. For each individual, mean values were obtained from 2 culture wells, each resulting from a distinct embryo, from at least 3 independent cultures. For NO measurement, enteric neurons were treated with pooled f-EVs from 15 control individuals or from 35 individuals with ASD, at a final concentration of 10^9^ particles/ml.

### Immunostaining and microscopy

#### Intestinal tissues

Segments of colon from treated mice were fixed in PBS containing 4% paraformaldehyde for 4 h at RT. Whole mounts of longitudinal muscle and myenteric plexus were obtained by microdissection and were permeabilized in PBS containg 0.5% Triton X-100 and 10% BSA for 2h at RT. Tissues were then incubated with human anti-Hu (1:1000, A21271, Molecular Probes) and rabbit anti-nNOS (1:500; ALX210529C100, Enzo Life Science) antibodies. After washing, tissues were incubated with appropriate AF 488- and Cy3-conjugated secondary antibodies for 2 hours at RT. Tissues were mounted with ProLong Gold Antifade Reagent.

#### Enteric neuron cultures

Cells were fixed in PBS containing 4% paraformaldehyde for 15 min at RT, permeabilized for 5 min in PBS containing 0.25% Triton X100 and incubated in PBS containing 10% BSA for 30 min. The cells were then incubated overnight at 4°C with the following primary antibodies diluted in PBS containing 3% BSA: rabbit anti-nNOS (1:500 ; ALX210529C100, Enzo Life Science), rabbit anti-synapsin-1 (1:1.000, D12G5, Cell Signaling), mouse anti-PSD95 (1:500,, Thermo Fisher Scientific), free fatty acid receptor 3 (FFAR3) After washing, the cells were incubated for 90 min at RT with the appropriate FP 488-, Cy3- or Alexa 568-conjugated secondary antibody diluted in PBS containing 3% BSA and were mounted in ProLong Gold Antifade Reagent.

#### Microscopy and quantification

Images were acquired with a digital camera (Hamamatsu C11440) coupled to a fluorescence microscope (Axio observer Zeiss) controlled via Zen software (Zeiss). The number of nNOS-positive neurons was counted in 8 ganglia for each colon and in 8 fields per well for neuron cultures from at least three independent cultures, and was normalized to the total number of Hu-positive neurons. To determine the number of synapsin 1 and PSD95 clusters, images were acquired with a x63 objective and a threshold was set with Fiji software to be two times higher than the underlying fluorescence intensity of the axonal processes. The number of synapsin 1 and PSD95 clusters were determined using the Fiji particle analysis tool with a cut off particle size of 0.2 µm^2^. The number of clusters was normalized to the axonal length.

### Ca^2+^ imaging

Neurons were loaded with Cal-520 AM dye (5 µM, ab171868, Abcam) in enteric glia-conditioned neurobasal medium for 60 min at 37°C and were then transferred to a temperature- and gas-controlled chamber maintained at 37°C and 5% CO2, mounted on a resonant scanning NIKON A1 Rsi confocal microscope (Nikon Instruments, France). High-frame-rate images (60 fps) were recorded with a x63 objective and the videos were acquired using the Nikon NIS-Elements software. Video analysis was performed using ImageJ software. The fluorescence intensity was determined in the region of interest (ROI) delimitating neuronal cell bodies. Data were expressed as changes in fluorescence intensity relative to the basal fluorescence level (F0), calculated as ΔF/Fo. For spontaneous activity, Fo was defined as the minimum fluorescence intensity over the entire recording period. For acute f-EV and pharmacological stimulation, Fo corresponded to the basal fluorescence intensity measured before stimulation. The analysis of Ca^2+^ transient amplitude and duration were performed using an Excel macro designed at the Plateforme MicroPicell (Nantes University). For acute treatments, neurons were stimulated with f-EVs (10^9^ particles/ml) derived from each individual or with acetylcholine (10 µM, A2661-25G, Sigma-Aldrich), serotonin (5-HT, H-9523, Sigma-Aldrich), or gamma-aminobutyric acid (GABA, 10 µM, Cat: A2129, Sigma-Aldrich). For acute treatment with pooled f-EVs, neurons were stimulated with either a mixture of control EVs derived from 15 control individuals or a mixture of ASD-EV derived from 35 individuals with ASD, at a final concentration of 10^9^ particles/ml. Treatments were performed in the absence or presence of the FFAR3 antagonist, β-hydroxybutyrate (5 mM) or of a mixture of butyrate (1.7 µM) and valerate (0.4 µM).

### Statistics

All data are presented as mean ± SEM (Standard Error of the Mean). Normality and equal variance were confirmed using the D’Agostino & Pearson, Anderson-Darling, Shapiro–Wilk and Kolmogorov-Smirnov tests, respectively. Comparisons of normally distributed data were conducted using t-tests with Welch’s correction or one-way ANOVAs, followed by post hoc Tukey’s tests. To assess the effects of two factors, a two-way ANOVA with Bonferroni’s multiple comparisons test was used. For non-parametric datasets, the Mann-Whitney test or the Kruskal-Wallis test, followed by Dunn’s multiple comparisons test, was used. Comparison test analyses were performed with GraphPad Prism software (GraphPad Software, Inc., La Jolla, CA, USA). Spearman’s rank correlation coefficient was employed for the correlation analyses using RStudio software (2025.5.1.513 version, Posit Software, PBC, Boston, MA) with the ‘hmisc’ package for data analysis, high-level graphics, missing value imputation, and advanced table-making. Significance was defined as p < 0.05. Each analysis consisted of two to three biological replicates from two to three independent cultures.

## RESULTS

### Metabolomic profiles differ between EVs from CTL individuals and ASD patients

Study participant demographics and clinical characteristics are summarized in Table 1. There were no significant difference in age or sex distribution between the two groups. Metagenomic analyses of the gut microbiome was performed in fecal samples from control individuals and patients with ASD. In total, 10 and 26 fecal samples were successfully sequenced from the CTL and ASD groups respectively. Alpha diversity, evaluated through richness and evenness, was similar between CTL and ASD individuals (Fig. 1A). Beta diversity, based on Bray-Curtis, Euclidean and Jaccard distances, was also similar between the two groups (Fig. 1B). Family-level microbiota profiling identified a selective depletion of members of the Lachnospiraceae family in ASD. Specifically, both *Lachnospiraceae H* and *Lachnospiraceae C* displayed reduced relative abundance compared with controls, although only the reduction in *Lachnospiraceae H* reached statistical significance. These findings point to a dysregulation of specific Lachnospiraceae taxa in ASD-associated gut microbial communities (Fig. 1C). Then, f-EVs were isolated from the same fecal samples from CTL individuals (f-EV-CTL) and individuals with ASD (f-EV-ASD). Transmission electron microscopy of isolated EVs showed the presence of bilayered spherical f-EVs from NT and ASD individuals (Fig. 1D). The size and number of f-EVs were analyzed by NTA. As shown in Fig. 1E, a distinct pattern of particle size distribution was found between NT and ASD individuals, with a bimodal distribution for f-EV-ASD compared to EV-NT (Fig. 1E). However, quantification of the particle size at the peak of frequency distribution showed no difference between NT and ASD individuals (Fig. 1E,F). The number of particles isolated from fecal samples was similar between CTL and ASD (Fig. 1G). The presence of bacteria-derived EVs was confirmed by the detection of lipopolysaccharide (LPS), measured at similar levels in f-EV-CTL and f-EV-ASD (Fig. 1H) as well as other bacterial protein markers, including LTA, OmpA and peptidoglycan (Fig. 1I). In addition, and as expected, markers for eukaryotic cell-derived EVs, CD63 and CD81 were also detected, suggesting the presence of combined microbiota- and host-derived EVs in fecal samples (Fig. 1J).

**Figure 1:**
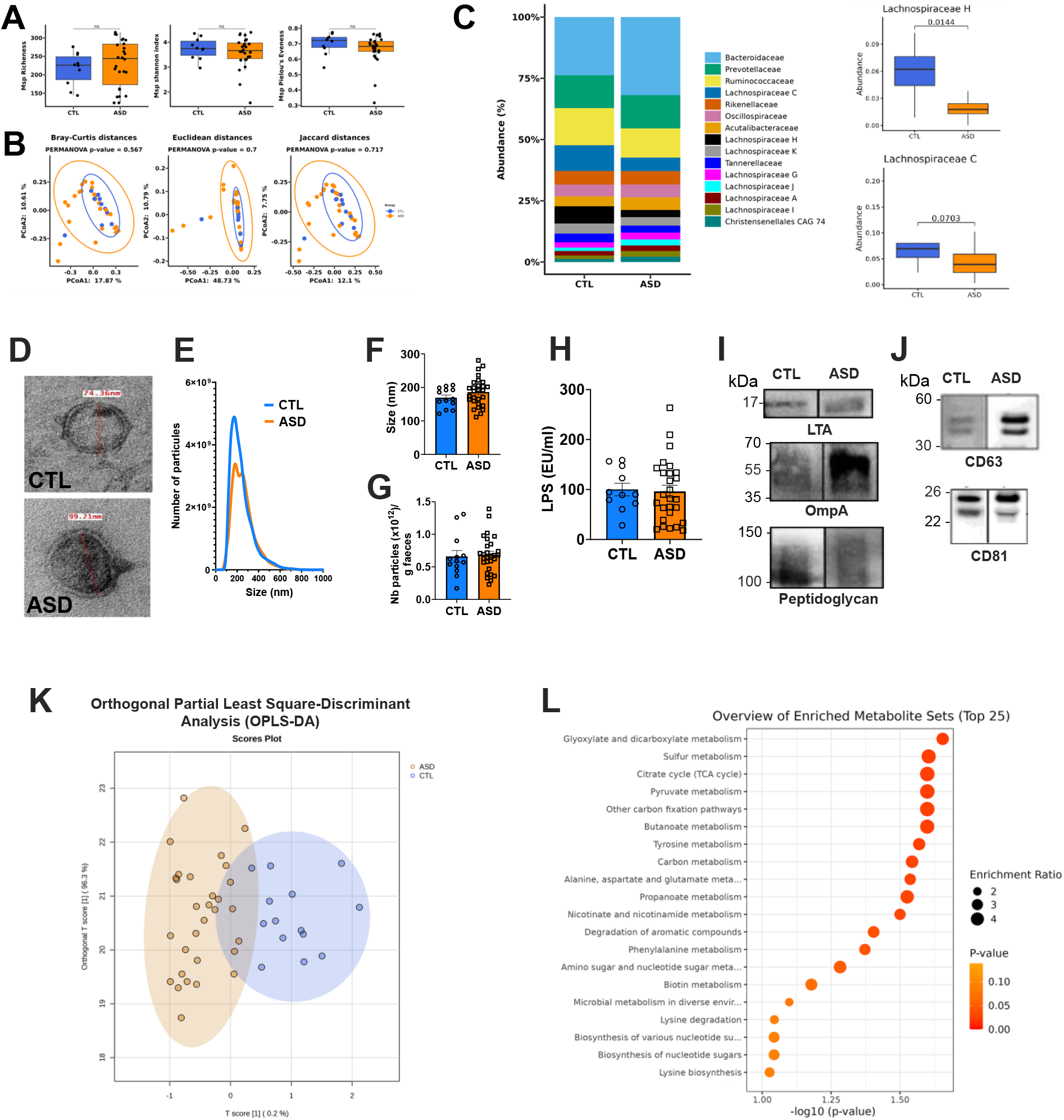
Gut microbiota composition and characterization of f-EVs from CTL and ASD individuals. (A) alpha microbiota diversity assessed by the richness, evenness (Pierlou’s index) and Shannon index. ns: non significant (CTL vs. ASD, p>0.05, Wilcoxon test). (B) Principal coord inate analysis (PCoA) of bacterial beta-diversity in CTL and ASD individuals generated by Bray–Curtis, Euclidean and Jaccard distances (CTL vs ASD, p>0.05, Permanova with 999 permutations). (C) Cumulative plot bar of the relative abundance of family bacterial taxon in of CTL and ASD individuals for the top 15 most abundant families. The boxplots highlight the reduced relative abundance for *Lachnospiraceae H* in the ASD group compared to CTL (p=0.014, non-parametric Wilcoxon tests). (D) Electronic microscopy images of f-EVs from a CTL individual and a patient with ASD. (E) Size distribution and mean (F) analyzed by nanoparticle tracking analysis (p= 0.16, Welch’s t-test). (G) Number of particles per gram of feces of f-EVs CTL: n=13, ASD: n=28; p= 0.4, Mann-Whitney test. (H) LPS concentration in f-EV-CTL (n=11) and f-EV-ASD (n=27) (p>0.05, Mann-Whitney test). (I, J) Representative western blot of f-EVs for bacterial markers (I) and EV markers (J). (K) Orthogonal partial least squares-Discriminant analysis based on 77metabolites identified in f-EV-CTL and f-EV-ASD (p =0.009, ANOVA). (L) Pathway enrichment analysis of f-EVs performed with MetaboAnalyst is represented in a dot-plot. Color intensity reflects statistical difference between f-EV-CTL and f-EV-ASD, and circle diameter shows pathway enrichment. Data are mean ± SEM. Statistical significance was determined using a Welch’s t-test or a Mann-Whitney test, depending on the data’s normality.

Next, using targeted metabolomics, we identified the metabolite content of f-EVs by mass spectrometry using the same particle number for each participant from the CTL (n=13) and ASD (n=28) groups. A total of 77 metabolites has been identified, including amino acids, aromatic compounds, lipids, nucleosides, and carbohydrates (Table S1). An Orthogonal Partial Least Squares Discriminant Analysis (OPLS-DA) based on these 77 metabolites, indicated that the CTL and the ASD groups were significantly distinguished suggesting that the metabolic composition of f-EV-ASD was different from f-EV-CTL (Fig. 1K). The KEGG pathway enrichment analysis indicated that differentially abundant metabolites were mainly related to the metabolism of glycoxylate/dicarboxylate, short chain fatty acids (SCFAs; butyrate, propionate), sulfur, pyruvate, tyrosine, carbon, alanine, aspartate, glutamate, and vitamine B family (nicotinate and nicotinamide) (Fig. 1L). Collectively, these findings identify altered SCFA signaling and neuroactive metabolite metabolism in f-EV-ASD as potential mechanistic drivers of the gastrointestinal and behavioral abnormalities associated with gut microbial dysbiosis.

### EVs from ASD patients alter social and anxiety-like behaviors and brain protein expression

#### ASD-derived f-EVs increase social interaction in recipient mice

To determine whether f-EVs from ASD patients influence behavior, mice receiving EVs from either control subjects (f-EV-CTL) or ASD patients (f-EV-ASD) were assessed for social, repetitive, and anxiety-related behaviors following the treatment period (Fig. 2A). Social behavior was first evaluated using the three-chamber sociability test. Both f-EV-CTL- and f-EV-ASD-treated mice spent significantly more time interacting with the chamber containing a social stimulus (mouse) than with the chamber containing a non-social stimulus (object), indicating intact sociability in both groups (Fig. 2B; f-EV-CTL: object vs. mouse, p = 0.0283; f-EV-ASD: object vs. mouse, p < 0.0001). Notably, f-EV-ASD-treated mice displayed a significantly higher sociability index than f-EV-CTL-treated mice (p = 0.0231), demonstrating enhanced social preference in response to ASD-derived f-EVs (Fig. 2B). This effect was consistently observed in both sexes, corresponding to male donor/male recipient and female donor/female recipient pairs (Supplementary Fig. S1A). Together, these findings indicate that ASD-derived f-EVs do not recapitulate the social interaction deficits classically associated with ASD. Rather, they promote social behavior in recipient mice, suggesting that microbiota-derived EVs selectively influence specific behavioral domains and highlighting the complexity of microbiota-host communication mechanisms in ASD.

**Figure 2.**
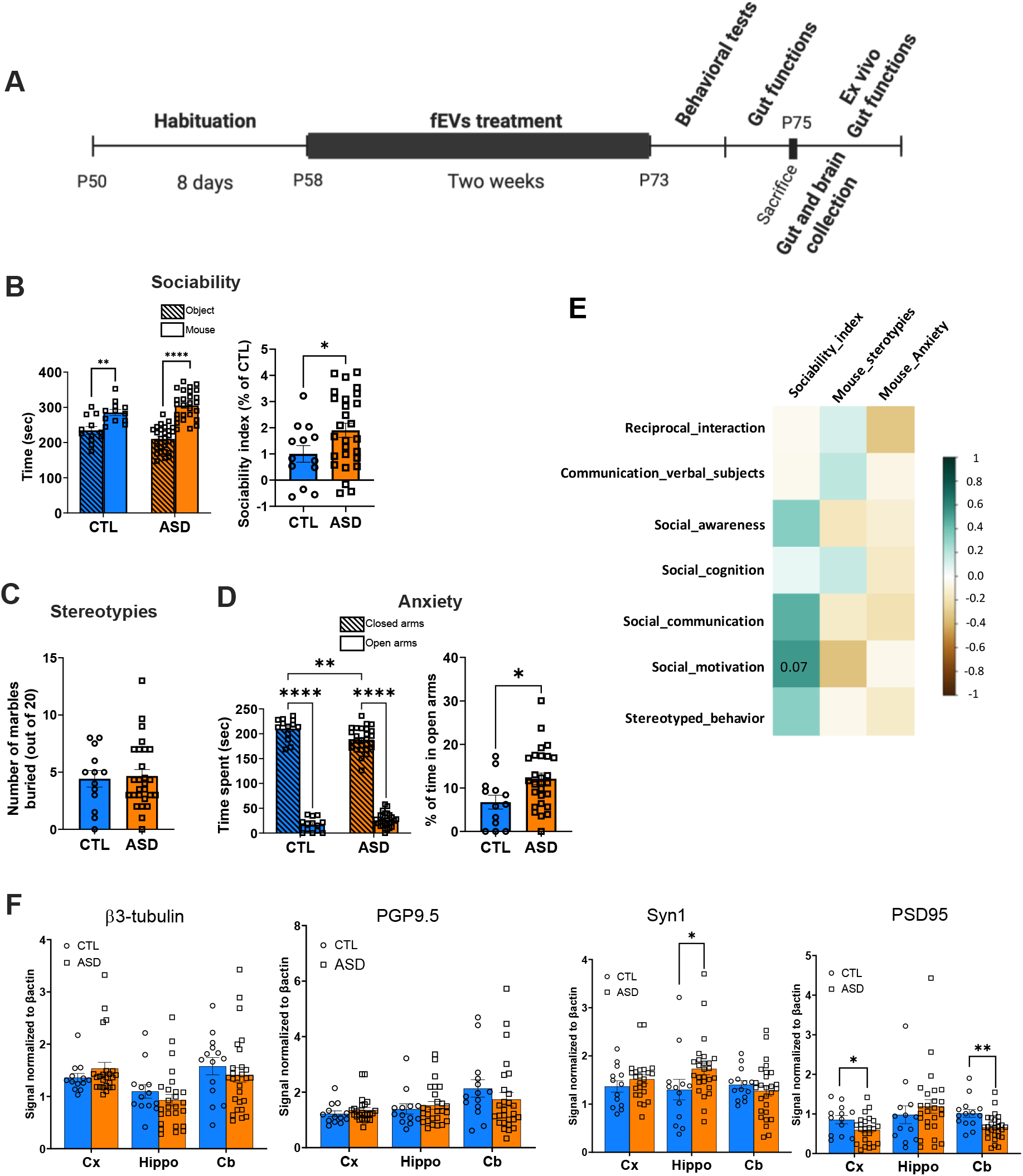
ASD patient-derived f-EVs alter social and anxiety-like behaviors and brain protein expression. (A) Experimental design of f-EV treatment and gut and brain analyses in mice. (B) Three-chamber sociability test: time spent by the mice in the chamber with unfamiliar sex- and age-matched mouse versus the chamber with an object as a non-social stimulus (**p=0.02 ****p<0.0001, two-way ANOVA with Bonferroni post hoc test, n=13 CTL, n=27 ASD). Right panel: sociability index (*p=0.04, Welch’s t-test). (C) Stereotypies assessed by the marble-burying test. The number of marbles buried by each mouse was counted over a 10-min period (p>0.05, Mann-Whitney test, n=13 CTL, n=27 ASD). (D) Elevated plus maze test, both f-EV-CTL and f-EV-ASD mice spend longer time in closed vs open arms (****p<0.0001 two-way ANOVA with Bonferroni post hoc test), but f-EV-ASD-treated mice spent longer time in open arms compared to f-EV-CTL-treated mice (**p=0.001). Right panel: percentage of time in open arms (*p=0.014, Welch’s t-test). (E) Spearman correlation matrix between f-EV-ASD-treated mice behavior scores and ADOS-2 scores of the corresponding patients. The correlation effect is indicated by a color gradient from brown (negative) to blue (positive). P values <0.1 are indicated (n=13 CTL, n=27 ASD). (F) Relative protein expression of the neuronal proteins beta-3-tubulin, PGP9.5, Synapsin-1 and PSD95 normalized to □-actin signal in the prefrontal cortex (Cx), hippocampus (Hippo), and cerebellum (Cb). Data are mean ± SEM. *p<0.05, **p<0.01, Mann-Whitney, n=13-14 CTL, n=24-28 ASD).

### ASD-derived f-EVs do not alter repetitive-like behavior

Repetitive and stereotyped behaviors were assessed using the marble-burying test. No significant difference in the number of buried marbles was detected between f-EV-CTL-and f-EV-ASD-treated mice (Fig. 2C; p = 0.86), indicating that ASD-derived f-EVs did not affect repetitive-like behavior under the experimental conditions tested.

#### ASD-derived f-EVs reduce anxiety-like behavior

Anxiety-related behavior was evaluated using the elevated plus maze. As expected, mice from both treatment groups spent significantly more time in the closed arms than in the open arms of the maze (Fig. 2D; p < 0.0001), reflecting the innate preference of mice for protected environments. However, f-EV-ASD-treated mice spent a greater proportion of time in the open arms compared with f-EV-CTL-treated mice (Fig. 2D; p = 0.0142), consistent with an anxiolytic-like effect of ASD-derived EVs. This anxiolytic effect was observed in both male and female mice, although it appeared more pronounced in females (Supplementary Fig. S1B).

Taken together, these data indicate that ASD-derived f-EVs selectively modulate behavioral responses by enhancing sociability and reducing anxiety-like behavior without affecting repetitive-like behaviors.

#### Behavioral effects of f-EV-ASD are associated with clinical social symptom severity

To investigate whether behavioral outcomes in recipient mice reflected clinical characteristics of the corresponding ASD donors, behavioral scores from f-EV-ASD-treated mice were correlated with clinical measures. A positive association was observed between the sociability index of recipient mice and the Social Motivation score of the corresponding patients on the ADOS-2 assessment. Although this relationship did not reach statistical significance, it showed a clear trend toward association (Spearman correlation, p = 0.07; Fig. 2E).

We next investigated the existence of possible brain alterations by assessing the expression levels of proteins involved in the structure and connectivity of the neuronal network. Western blot analyses were performed in three brain regions relevant to the clinical phenotype of ASD, the frontal cortex, hippocampus and cerebellum^32^. The proteins examined included βIII-tubulin, a microtubule-associated protein highly enriched in the axonal processes and commonly used as an indicator of axonal arborization; PGP9.5, a cytoplasmic hydrolase abundantly expressed in neurons and widely used as a marker of neuronal density; synapsin 1, a presynaptic protein used as a marker of presynaptic terminals; PSD95, a scaffolding postsynaptic protein used as a marker of postsynaptic sites. No differences in the expression of βIII-tubulin and PGP9.5 were observed between f-EV-CTL- and f-EV-ASD-treated mice while f-EV-ASD-treated mice showed an increased expression of synapsin-1 in the hippocampus (Fig. 2F; p=0.01) and reduced PSD95 levels in the prefrontal cortex and cerebellum compared to f-EV-CTL treated mice (p=0.04 and 0.007 respectively; Fig. 2F and Suppl Fig. S1C). These findings suggest that ASD-derived f-EVs selectively modulate synaptic organization and function in a brain region-specific manner, without affecting overall neuronal abundance.

### Ileal and colonic permeability are unaffected by f-EV-ASD treatment

The potential impact of EVs treatment on gut functions was first studied on the intestinal epithelial barrier permeability. The paracellular permeability, assessed *in vivo* by measuring plasmatic sulfonic acid (0.4 kDa) and dextran (4 kDa) concentration after gavage was unaffected by f-EV-ASD treatment compared to controls (Fig. 3A) as well as the transcellular permeability measured by HRP (44 kDa) (Fig. 3A). To further characterize barrier permeability in specific gut regions, the ileum and the colon were collected to proceed for *ex vivo* measurement of sulfonic acid, dextran and HRP flux. No effect of f-EV-ASD was observed for any of the tested molecules, in either the ileum or the colon (Fig. 3B).

**Figure 3.**
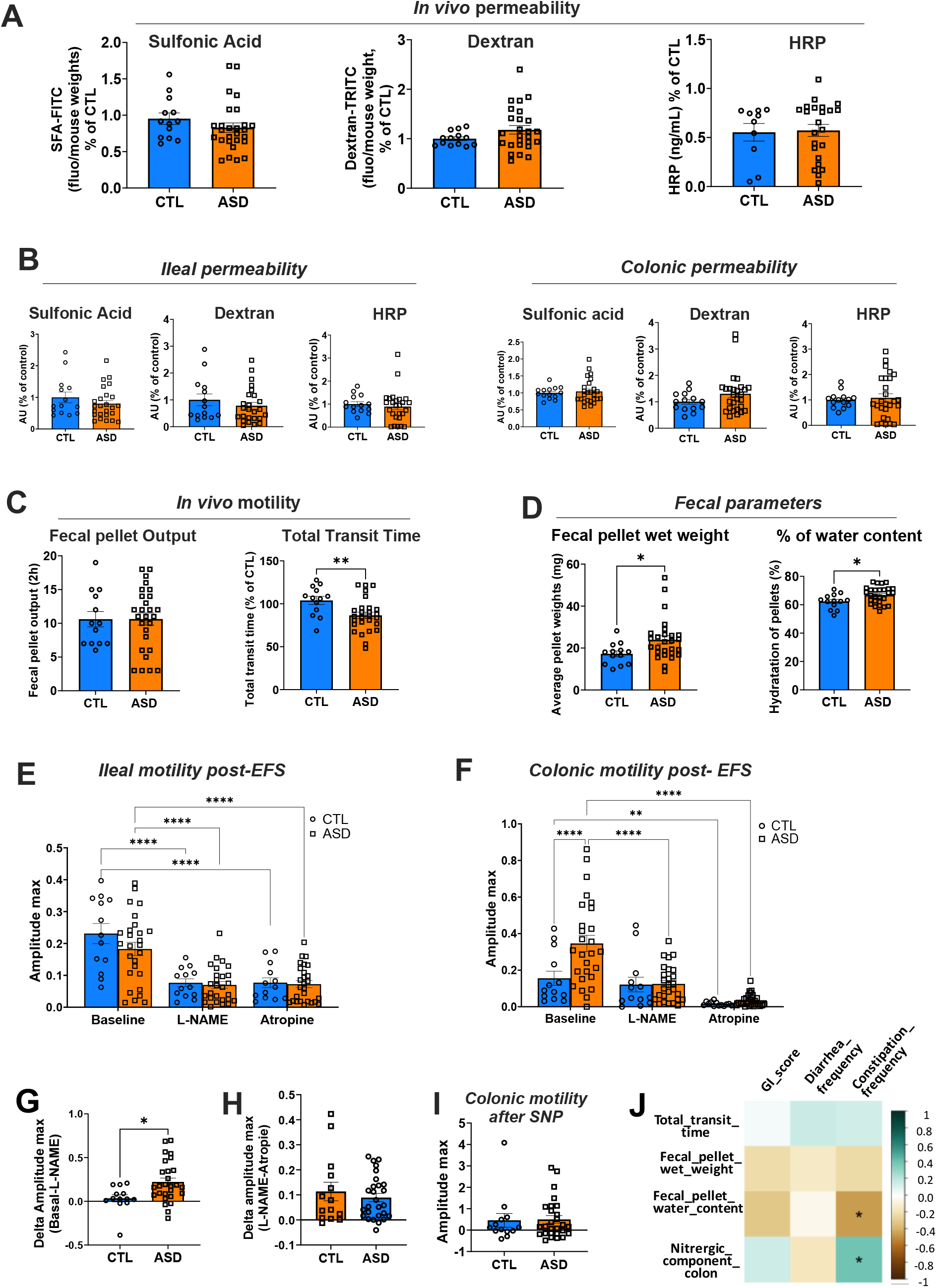
f-EV-ASD induce no change in intestinal barrier permeability but modulates colonic motility by enhancing the NO-sensitive component. (A) *In vivo* permeability for sulfonic acid, dextran and HRP in mice treated with f-EV-CTL or f-EV-ASD. Data are mean ± SEM (p>0.05, Mann-Whitney test). (B) E*x vivo* permeability for sulfonic acid, dextran and HRP in colon and ileal segment of mice treated with f-EV-CTL or f-EV-ASD (p>0.05, Mann-Whitney test). (C) *In vivo* GI motility assessed by fecal pellet output over a 2-h period and by the total transit time of carmine red dye in mice treated with f-EV-CTL or f-EV-ASD (**p=0.008, Mann-Whitney test). (D) Fecal parameters of mice treated with f-EV-CTL or f-EV-ASD assessed by fecal pellet wet weight and the percentage of water content (*p=0.02, Mann-Whitney test). (E) Maximum amplitude of the post-EFS response of ileal segments and colon segments (F) from f-EV-CTL- and f-EV-ASD-treated mice in basal condition and after sequential addition of L-NAME (50 µM) and atropine (1 µM). ****p<0.0001, **p<0.01, 2-way ANOVA repeated measure with Bonferroni post hoc test, n=13 CTL, n=27 ASD. (G) The L-NAME-sensitive maximum amplitude (difference between maximum amplitude in basal condition and in the presence of L-Name) in colon segments from f-EV-CTL- and f-EV-ASD-treated mice. *p<0.05, Mann-Whitney test. (H) The atropine-sensitive maximum amplitude (difference between maximum amp litude in the presence of L-NAME and in the presence of L-NAME + atropine) in colon segments from f-EV-CTL- and f-EV-ASD-treated mice (p>0.05, Mann-Whitney test). (I) Maximum amplitude of the response of colon segments upon SNP application (p>0.05, Mann-Whitney test). (J) Spearman correlation matrix between the GI functions of f-EV-ASD-treated mice and the GI conditions reported in the corresponding patients. The correlation effect is indicated by a color gradient from brown (negative) to blue (positive). Significant associations are marked with asterisks. *p<0.05, Spearman’s rank correlation analysis.

### Treatment with f-EV-ASD decreases gastrointestinal transit time and affects colonic motility through a higher contribution of the NO-sensitive component

Gastrointestinal motility was studied *in vivo* by assessing the colonic propulsive motor functions through FPO measurements. We found a comparable FPO values between f-EV-CTL- and f-EV-ASD-treated mice (Fig. 3C). However, a shorter total transit time was observed in mice treated with f-EV-ASD compared to those treated with f-EV-CTL (Fig. 4C; p=0.008), in both males and females, although the effect was more pronounced in males (Suppl Fig. S2A). Feces analyses indicated an increased fecal pellet wet weight and water content in mice treated with f-EV-ASD compared to the control group (Fig. 3D; p=0.02). Further characterization of gut motility was conducted *ex vivo* to assess the neuromuscular response of ileal and colonic segments placed in an organ chamber and submitted to electrical field stimulation (EFS). Measurements were performed 60 sec after the EFS termination (post-EFS). In the ileum, the amplitude of the post-EFS-induced contractile response was similar for the f-EV-ASD and f-EV-CTL group (Fig. 3E; baseline values). Following the addition of the nitric oxide synthase inhibitor (L-NAME), post-EFS-induced contractile activity was similarly reduced in the f-EV-CTL and f-EV-ASD groups (Fig. 3E). Similarly, the addition of atropine, a muscarinic receptor antagonist, equally decreased the post-EFS-induced contractile activity in the two groups (Fig. 3E). In the colon, under basal condition, a greater amplitude of post-EFS-induced response was observed for f-EV-ASD compared to f-EV-CTL groups (Fig. 3F). After the addition of L-NAME, the amplitude of the post-EFS-induced response was significantly reduced in the f-EV-ASD group (Fig. 3F). Following the addition of atropine, post-EFS-induced contractile response was reduced in both the f-EV-CTL and f-EV-ASD groups (Fig. 3F). Comparison between the two groups for the L-NAME-sensitive component in response to EFS indicated a greater sensitivity to the NOS blockade in the colon segments from f-EV-ASD-treated mice than those from f-EV-CTL-treated mice (Fig. 3G, p=0.02). No difference was observed regarding the atropine-sensitive response between the two groups (Fig. 3H). Interstingly, the same results were obtained in male and female mice (Suppl Fig. S2B). We then assessed in the colon whether muscle sensitivity to NO was affected by f-EV-ASD treatment by investigating the muscle response of colonic segments to the NO donor sodium nitroprusside (SNP, 100 µM). No difference in the colon was found between the two groups, suggesting that the sensitivity of the muscles to NO was not modified by the f-EV-ASD treatment (Fig. 3I). These results suggest that f-EV-ASD treatment modified colonic motility, at least in part, through remodeling of the nitrergic component of the ENS. Correlation analysis between the gastrointestinal conditions reported in patients and the gastrointestinal functions assessed in mice revealed that the frequency of constipation in patients was negatively associated with fecal water content and positively associated with the extent of the nitrergic component of colonic motility in f-EV-ASD-treated mice. (Spearman’s p<0.05; Fig. 3J). The pharmacological analyses indicate that f-EV-ASD treatment selectively increases the nitrergic contribution to colonic neuromuscular transmission without affecting cholinergic signaling. The greater sensitivity to NOS inhibition, together with the unchanged responses to atropine and the NO donor, SNP, suggests that f-EV-ASD primarily remodels enteric nitrergic neuronal pathways rather than smooth muscle responsiveness to nitric oxide or cholinergic neurotransmission. This enhanced nitrergic tone may contribute to the altered colonic motility and constipation-associated phenotype observed in f-EV-ASD-treated mice. In addition, because nitric oxide signaling within the intestinal submucosa can arise from functional interactions between enteric neurons and the microvascular endothelium, a contribution of neuronal-endothelial crosstalk cannot be excluded. Indeed, neurotransmitters released from submucosal neurons have been shown to stimulate endothelial NO release and regulate intestinal vascular function (Acetylcholine released from guinea-pig submucosal neurones dilates arterioles by releasing nitric oxide from endothelium (R Andriantsitohaina, A Surprenant, J Physiol. 1992:453:493-502), suggesting that f-EV-ASD-induced alterations in NO signaling may involve coordinated actions of enteric nitrergic neurons and endothelial cells.

**Figure 4:**
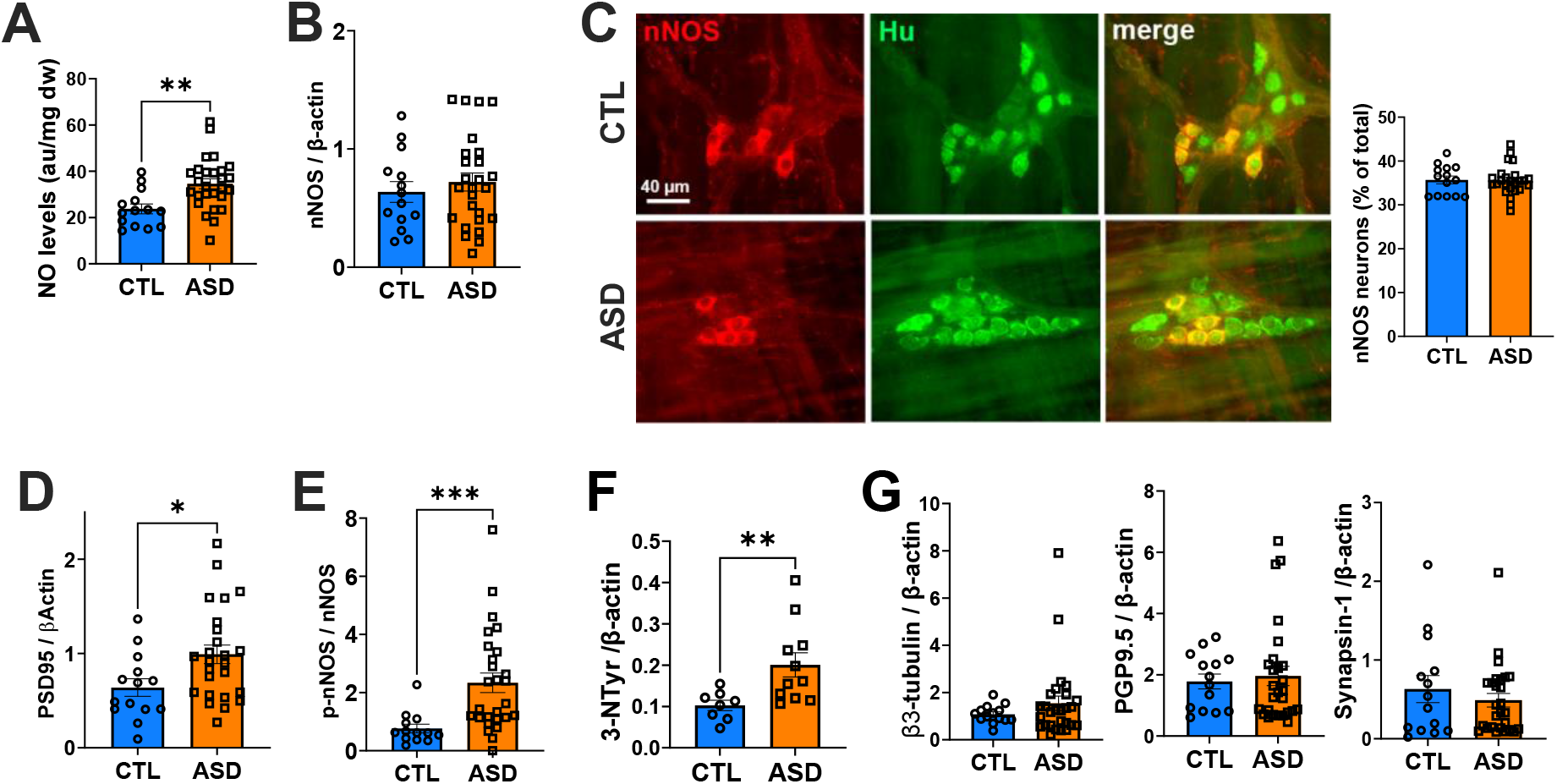
Increased NO levels, nNOS activation and protein tyrosine nitration in the colon of mice treated with f-EV-ASD. (A) NO concentration in the colon of f-EV-CTL- and f-EV-ASD-treated mice (**p<0.01, Student’s t test, n=14 CTL, n=27 ASD). (B) Protein expression levels of nNOS normalized to β -actin signal in the colon of mice treated with f-EV-CTL or f-EV-ASD. (p>0.05, Mann-Whitney test). (C) Double immunolabeling for nNOS (red) and HuC/D (neuronal marker, green) in the colonic myenteric plexus of mice treated with f-EV-CTL or f-EV-ASD. Right panel: quantitative analysis for the proportion of nNOS-positive neurons (p>0.05, Mann–Whitney test). (D,E,F) Protein expression levels normalized to □-actin signal in the colon of mice treated with f-EV-CTL or f-EV-ASD for PSD95 (D), p-nNOS at Ser1412 (E) and 3-NTyr as a marker of protein tyrosine nitration (F), *p=0.01, **p=0.008, ***p=0.0002, Mann-Whitney test. (G) Relative protein expression of the neuronal proteins beta-3-tubulin, PGP9.5 and synapsin-1 normalized to β-actin signal in the colon of mice treated with f-EV-CTL or f-EV-ASD (p>0.05, Mann-Whitney test, n=13-14 CTL, n=27 ASD).

### Treatment with f-EV-ASD enhances NO concentration, nNOS activation and protein tyrosine nitration in the colon

To further explore the changes of the NO pathway induced by the f-EV-ASD treatment, we next assessed in colonic tissues the NO levels and the expression of the neuronal NO-producing enzyme nNOS. We found an increased NO concentration in the colon of f-EV-ASD-treated mice as compared to f-EV-CTL-treated mice (Fig. 4A; p=0.001), in both male and female mice (Suppl Fig. S2C), while nNOS expression level and number of nNOS-expressing neurons were unaffected (Fig. 2B, C). Notably, nNOS activity can be markedly regulated independently of changes in its expression levels through its interaction with the scaffolding protein PSD95, which recruits and anchors nNOS to specialized plasma membrane microdomains and promotes its functional activation^33–35^. Consistent with enhanced nitrergic signaling, f-EV-ASD-treated mice exhibited increased colonic PSD95 expression compared with f-EV-CTL-treated mice (Fig. 4D; p = 0.01), together with an increased proportion of the activated Ser1412-phosphorylated form of nNOS (P-nNOS) (Fig. 4E). These findings indicate that f-EV-ASD enhances nNOS activation rather than nNOS expression per se. The increases in PSD95 and P-nNOS were observed in both male and female mice (Suppl. Fig. S2D). The increase of NO levels and nNOS activation could impact NO-mediated cell signaling, in particular those involved in protein tyrosine nitration previously shown to be elevated in the blood of children with ASD^36^. Protein tyrosine nitration corresponds to posttranslational modification of proteins through tyrosine nitrosative reactions achieved by NO-derived peroxynitrite, which eventually affect protein function, activity or protein interaction specificity^37^. We found that colon from mice treated with f-EV-ASD showed a higher level of protein tyrosine nitration than those treated with f-EV-CTL (Fig. 4F; p=0.008). Finally, we assessed the expression of neuronal markers indicative of neuronal network integrity and connectivity. Western blot analyses of colonic samples collected from f-EV-CTL- and f-EV-ASD-treated mice showed no differences between the two groups in the expression of β3-tubulin, PGP9.5 and synapsin 1 (Fig. 4G). Together, f-EV-ASD enhances the activity of existing nitrergic neurons, resulting in greater NO production and downstream nitrosative signaling, rather than inducing structural changes in the nitrergic neuronal population.

### f-EV-ASD enhance nNOS expression and NO levels in enteric neurons

Our findings showing that f-EV-ASD modulate the NO pathway in mice colon, in particular the phosphorylation of the neuronal form of the enzyme NOS, prompted us to study whether f-EV-ASD could directly impact enteric neurons and modulate neuronal NO pathway. Preliminary experiments were performed using different concentrations of f-EV-CTL and f-EV-ASD, which led to the selection of a final working concentration of 10⁹ particles/ml (Suppl Fig. S3A). Primary cultures of rat enteric neurons were treated for 2 days with f-EV-CTL or f-EV-ASD (10^9^ particles/ml) and the expression of nNOS was assessed by Western blot and immunofluorescence. Treatment with f-EV-ASD increased nNOS protein expression in enteric neurons (Fig. 5A, B). Consistently, NO levels were significantly higher in the culture medium from f-EV-ASD-treated neurons compared to f-EV-CTL-treated neurons (Fig. 5C). In addition, similar to the *in vivo* studies, PSD95 protein expression was increased following f-EV-ASD treatment compared to controls (Fig. 5D). Interstingly, we observed an increased number of PSD95 clusters on nitrergic neurons (Fig. 5E) and an increased PSD95 cluster density at synaptic sites, as indicated by a higher number of PSD95 clusters colocalizing with the synaptic marker synapsin 1 (Fig. 5F). Taken together, these findings suggest that EVs can directly act on enteric neurons and that EVs isolated from patients with ASD can promote nNOS expression and NO production.

**Figure 5:**
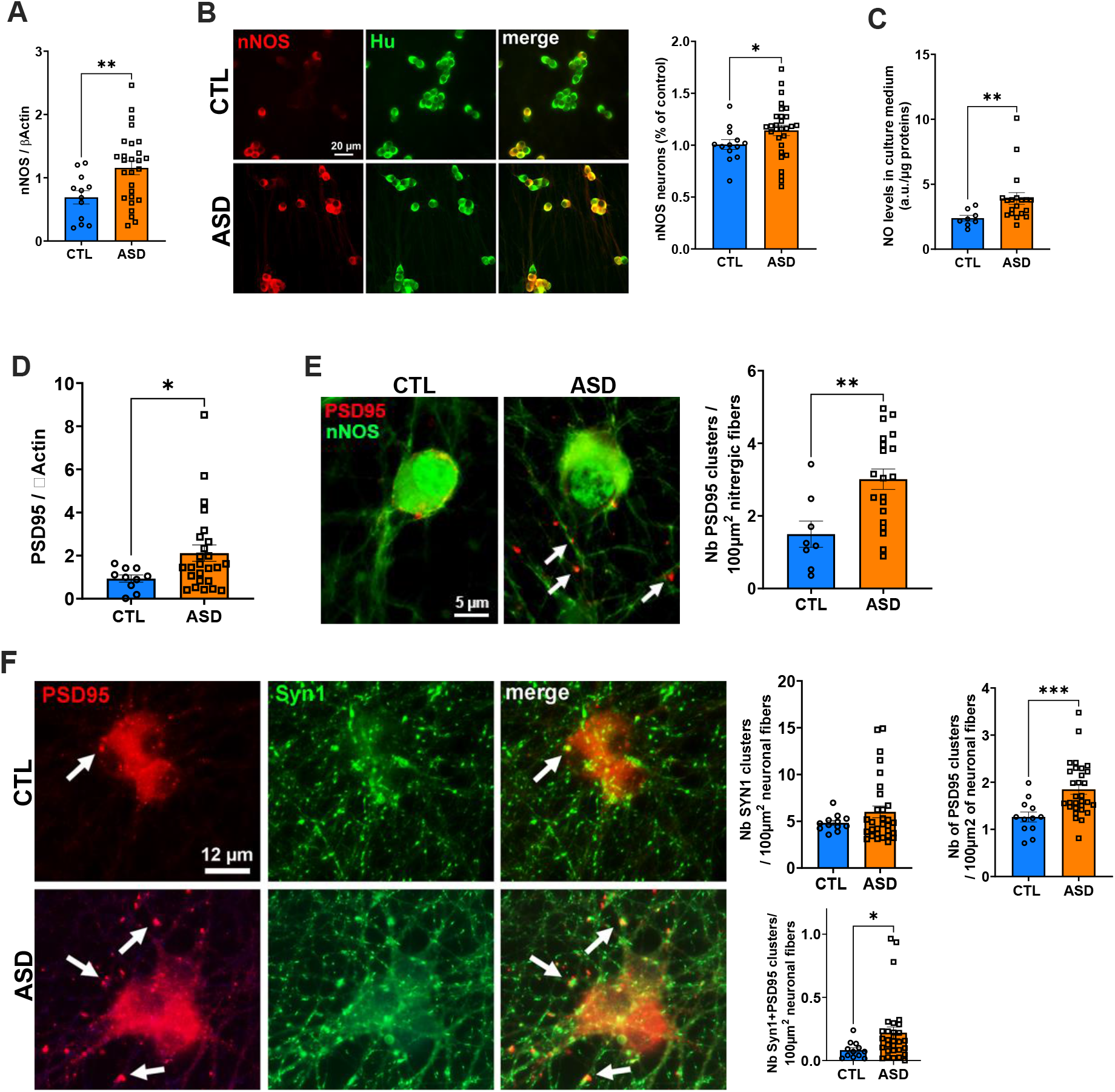
f-EV-ASD enhance nNOS expression, NO levels and synaptic PSD95 cluster number in enteric neurons. (A) nNOS protein expression normalized to beta-actin signal in cultured enteric neurons treated for 48h with f-EV-CTL or f-EV-ASD (**p=0.04, Mann-Whitney test, n=8 CTL, n=19 ASD). (B) Double immunolabeling for nNOS (red) and Hu (green) in enteric neurons after 48h of f-EV treatment. Right panel: quantitative analysis for the proportion of nNOS-positive neurons (*p=0.04, Mann–Whitney test). (C) NO concentration in the culture medium of enteric neurons treated for 48h with EVs (**p=0.004, Welch’s t test). (D) Protein expression levels of PSD95 normalized to β-actin signal in enteric neurons treated for 48h with f-EVs (*p=0.04, Mann-Whitney test). (E) Double immunolabeling for PSD95 (red, arrows) and n-NOS (green) in enteric neurons treated for 48h with f-EVs. Right panel: quantitative analysis for the number of PSD95 clusters in nNOS-positive neurons (**p=0.06, Welch’s t-test). (F) Double immunolabeling for PSD95 (red, arrows) and synapsin 1 (green) in enteric neurons treated for 48h with f-EVs. Right panels: quantitative analyses for the number of synapsin, PSD95 and colocalized synapsin-PSD95 clusters (*p=0.02, ***p=0.0007, Mann-Whitney test). (A,B,D,E,F): each data point represents the mean of two independent cultures well treated with EVs from n=10-12 CTL and n=25-30 ASD from three independent cultures. C: Each data point represents one culture well from a single embryo treated with pooled f-EVs from 15 control individuals or 35 patients with ASD at a final concentration of 10^9^ particles/ml and performed in three independent cultures.

### The f-EV-ASD alter spontaneous Ca^2+^ transients in enteric neurons and induce a sustained elevation of intracellular Ca^2+^

A valuable approach to assess the physiological status of neurons is to analyze spontenous intracellular Ca^2+^ transients over time. In addition, because nNOS activation and NO production in neurons are critically dependent on intracellular Ca^2+^ dynamics^38^, alterations in spontaneous Ca^2+^ dynamics may influence NO production. To study whether EV treatments modulate spontaneous intracellular Ca^2+^ dynamics, cultured neurons were treated with f-EV-CTL or f-EV-ASD for 2 days, and spontaneous Ca^2+^ transients were recorder in neuronal cell bodies for 2 min using live Ca^2+^ imaging. Quantitative analysis of spontaneous Ca^2+^ transients revealed a slight reduction in the peak amplitude, together with a prolonged peak duration, in f-EV-ASD-trated neurons compared with f-EV-CTL-treated neurons (Fig. 6A). Consistent with these findings, the decay slope was significantly decreased in response to EV treatment, irrespective of donor sex, indicating that EVs from both male and female individuals exerted similar effects. (Suppl Fig. S3B). We next investigated whether f-EV-ASD treatment might influence the response of enteric neurons to evoked stimulation by key neurotransmitters. Enteric neurons exposed to f-EV-CTL or f-EV-ASD for 2 days were challenged with acetylcholine (ACh), serotonin (5-HT) and GABA. No difference was observed between f-EV-CTL or f-EV-ASD groups on the amplitude and duration of the neurotransmitter-induced intracellular Ca^2+^ peaks (Fig. 6B, C, D).

**Figure 6:**
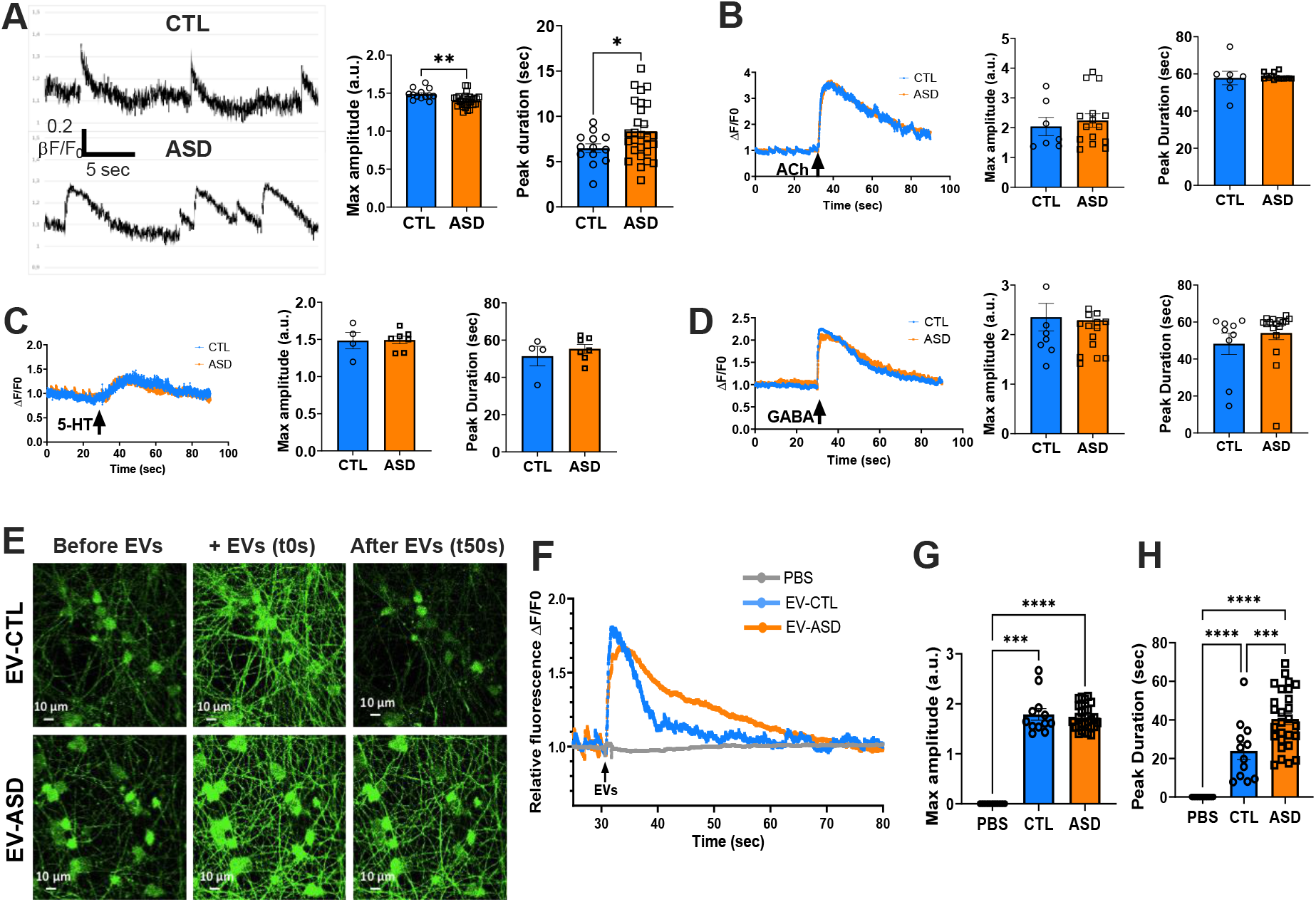
f-EV-ASD alter spontaneous Ca^2+^ transients in enteric neurons and induce a sustained elevation of intracellular Ca^2+^. (A) Representative traces of spontaneous Ca^2+^ transients in enteric neurons treated for 48h with f-EV-CTL or f-EV-ASD. Right panels: quantification of the maximum amplitude and duration of Ca^2+^ peaks. Each point corresponds to one individual (n=15 CTL, n=30 ASD) corresponding to the mean of 10-15 neurons recorded in one culture well from three independent cultures. *p=0.02, **p=0.003, Welch’s t-test. (B, C, D) Traces and quantification of evoked Ca^2+^ response of enteric neurons pretreated for 48h with pooled f-EV-CTL from 15 individuals or pooled f-EV-ASD from 30 patients and stimulated with Ach (10 µM, B), 5-HT (10 µM, C) or GABA (10 µM, D) (p>0.05, Mann-Whitney test). (E) Representative Cal-250 fluorescence images acquired before, during, and 50 s after the addition of f-EV-CTL or f-EV-ASD. (F) Representative traces of Ca^2+^ response evoked by vehicle (PBS), f-EV-CTL or f-EV-ASD. (G,H) Quantification of the evoked Ca^2+^ peaks for the maximum amplitude (G) and peak duration (H). Each point represents to one individual (n=12 CTL, n=24 ASD) corresponding to the mean of 10-15 neurons recorded in two culture wells from two independent cultures (***p<0.001, ****p<0.0001, ANOVA).

Next, we investigated whether EVs could directly elicit an acute intracellular Ca^2+^ response in enteric neurons. Application of f-EV-CTL and f-EV-ASD on cultured enteric neurons induced a rapid and transient increase in the intracellular Ca^2+^ response in similar proportion of enteric neurons (86.5±2.8% and 89.8±1.9% respectively; p=0.34). However, the profile of Ca^2+^ responses differed between the two groups, with a more sustained Ca^2+^ elevation observed for f-EV-ASD than f-EV-CTL (Fig. 6E, F). Acute application of f-EV-ASD induced no change in the peak amplitude of the Ca²⁺ response but resulted in a prolonged duration of Ca²⁺ elevation compared with f-EV-CTL (Fig. 6G, H). Consistent with this finding, the decay slope was significantly reduced, an effect that was more marked in mice treated with EVs from male donors than in those receiving EVs from female donors (Suppl Fig. S3C). These findings establish EVs as direct modulators of enteric neuronal activity and show that f-EV-ASD potentiates and prolongs intracellular Ca²⁺ signaling relative to f-EV-CTL.

### Reduced levels of butyrate and valerate in f-EV-ASD regulate Ca^2+^ response through the FFAR3-dependent signaling pathway

Given that our pathway enrichment analysis identified dysregulated SCFA metabolism in f-EV-ASD compared to f-EV-CTL, we quantified SCFA levels to identify EV-associated bacterial mediators that may be involved in the effects induced by f-EV-ASD on the colon and enteric neurons (Fig. 1L). We found that f-EV-ASD exhibited reduced levels of butyrate, valerate and isovalerate compared with f-EV-CTL (Fig. 7A). Previous studies have shown that SCFAs, especially butyrate and valerate, regulate colonic activity by activating the free fatty acid receptor 3 (FFAR3), a G protein-coupled receptor^39^. Considering that expression of G protein-coupled receptors is highly regulated by the level of ligand-induced activation^40^, we analyzed the protein expression of FFAR3 in mouse colon treated with f-EV-CTL and f-EV-ASD. We found a reduced FFAR3 expression in the colon of f-EV-ASD-treated mice compared with f-EV-CTL-treated mice (Fig. 7B). Next, we investigated whether SCFA-mediated signaling may be involved in EV-induced enteric neuronal Ca^2+^ transients. First, the expression of FFAR3 in cultured enteric neurons was examined by immunofluorescence. FFAR3 immunoreactivity was associated to both nitregic and non-nitregic enteric neurons and was found to be distributed at discrete puncta throughout neuronal cell bodies and axonal fibers (Fig. 7C). Then, enteric neurons were monitored for Ca^2+^ imaging upon acute f-EV-CTL application in the presence of the FFAR3 antagonist, β-hydroxybutyrate. FFAR3 receptor blockade resulted in a more sustained elevation of intracellular Ca^2+^ following f-EV-CTL stimulation, as evidenced by an increased peak duration (Fig. 7D). Conversely, enteric neuron stimulation with f-EV-ASD supplemented with butyrate and valerate attenuated the Ca^2+^ response, leading to a shorter Ca^2+^ peak duration compared with f-EV-ASD alone (Fig. 7E). Collectively, these results identify FFAR3-dependent SCFA signaling as a key regulator of enteric neuronal Ca²⁺ dynamics and suggest that reduced butyrate and valerate levels in f-EV-ASD contribute to the sustained Ca²⁺ response induced by ASD-derived EVs.

**Figure 7:**
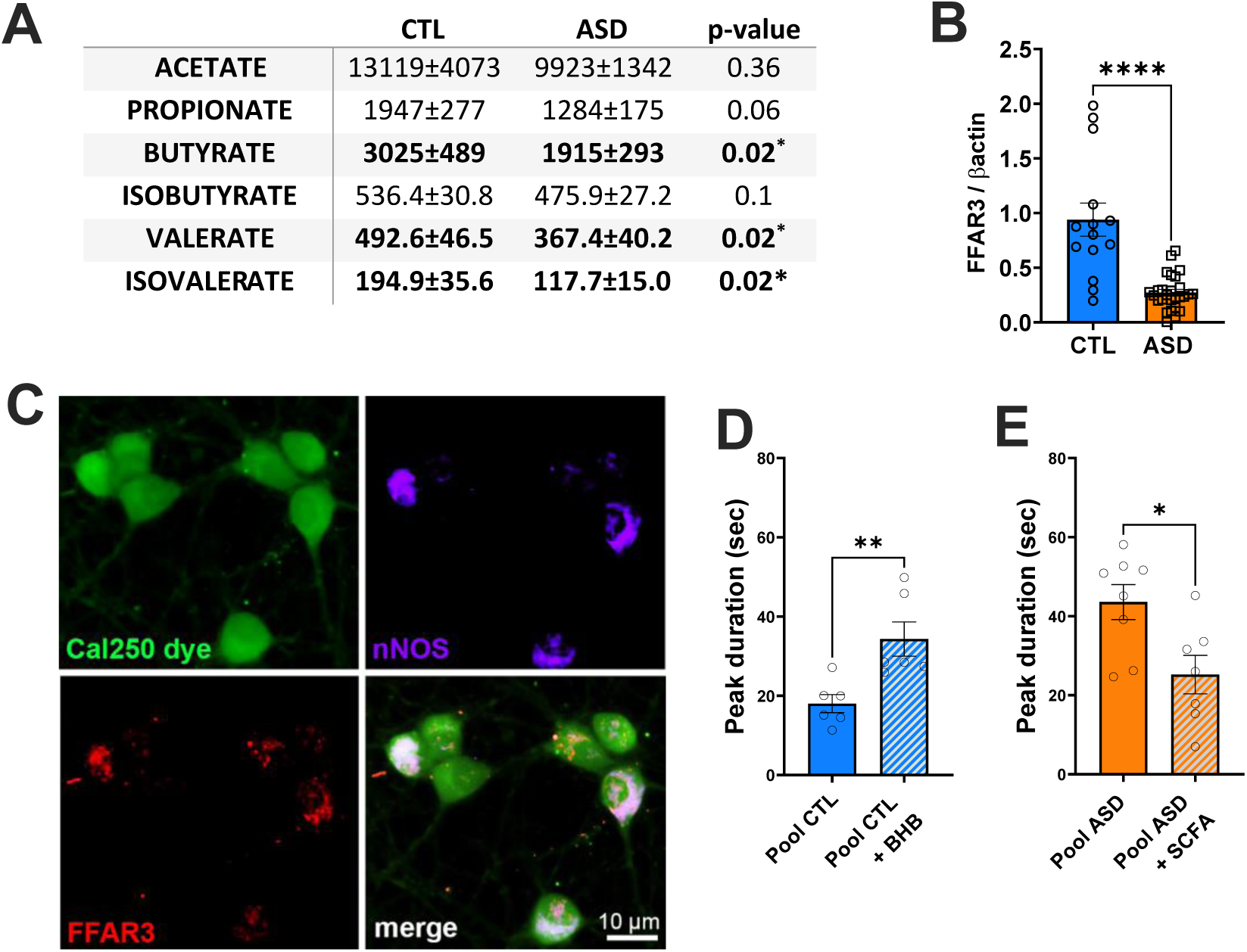
Reduced levels of SCFAs in f-EV-ASD and role of butyrate and valerate in EV-mediated Ca^2+^ response in enteric neurons. (A) SCFAs concentration in f-EV-CTL (n=15) and f-EV-ASD (n=30). Data are in nM and represent mean ± SEM. *p<0.05, Mann-Whitney test, n=13 CTL, n=24 ASD). (B) FFAR3 protein expression normalized to β-actin signal in the colon of mice treated with f-EV-CTL or f-EV-ASD (****p<0.0001, Mann-Whitney test, n=14 CTL, n=27 ASD). (C) Double immunolabeling for nNOS (purple) and FFAR3 (red) in enteric neurons. Neurons are visualized with Cal-250 dye (green). (D) Peak duration of Ca^2+^ transients evoked by f-EV-CTL in the presence or absence of the FFAR3 antagonist β-hydroxybutyrate (BHB, 5 mM). Each data point represents one culture well treated with pooled f-EV-CTL from 15 control individuals at a final concentration of 10^9^ particles/ml and performed in three independent cultures (**p=0.004, Mann-Whitney test, n=6). (E) Peak duration of Ca^2+^ transients evoked by f-EV-ASD in the presence or absence of 1.2 µM butyrate and 0.2 µM valerate. Each data point represents one culture well treated with pooled f-EV-ASD from 30 individuals with ASD at a final concentration of 10^9^ particles/ml and performed in three independent cultures (*p=0.02, Mann-Whitney test, n=7-8).

Correlation analysis between SCFA concentrations in f-EV-ASD and the clinical ADOS-2 score of the corresponding patients revealed a negative association between propionate, butyrate, isobutyrate, valerate, and isolavalerate concentrations and the communication score, as well as between isobutyrate concentration and reciprocal interaction (Spearman’s p<0.05; Fig. 8). Regarding the behavior of mice treated with f-EV-ASD, propionate and butyrate concentration showed a negative correlation with the sociability index, although this association did not reach statistical significance (Spearman’s p=0.08-0.09; Fig. 8). Correlation analysis between SCFA concentrations in f-EV-ASD and the clinical gastrointestinal symptoms of the corresponding patients indicated a negative association between isobutyrate and the diarrhea frequency, as well as between isovalerate and constipation frequency (Spearman’s p<0.05; Fig. 8C). Regarding the gastrointestinal phenotype of mice treated with f-EV-ASD, a positive association was found between the f-EV concentration of propionate and the total transit time, although this association did not reach statistical significance (Spearman’s p=0.05; Fig. 8D).

**Figure 8.**
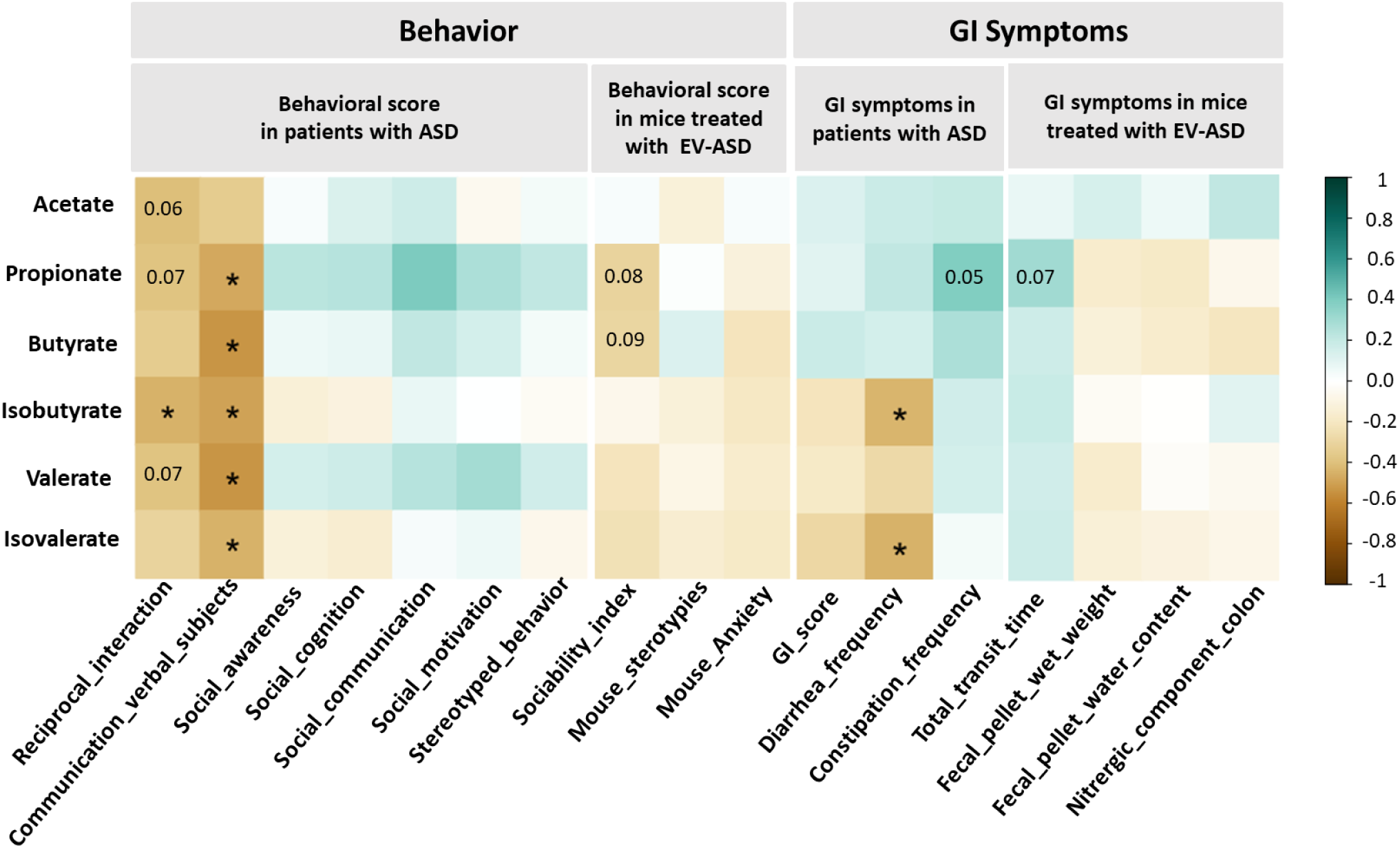
Negative association between SCFA levels in f-EV-ASD and social behavior and GI symptoms in patients with ASD. Spearman correlation matrix between the SCFA concentration in f-EVs from patients with ASD and behavior or GI symptoms in the corresponding patient with ASD or mouse treated with f-EV-ASD. The correlation effect is indicated by a color gradient from brown (negative) to blue (positive). Significant associations are marked with asterisks and p-values are reported when 0.05 < p < 0.1. *p<0.05, Spearman’s rank correlation analysis.

## DISCUSSION

In this study, we demonstrate that f-EVs from individuals with ASD possess a distinct molecular signature and are sufficient to induce both behavioral and gastrointestinal alterations in recipient mice. Specifically, f-EV-ASD reduced gastrointestinal transit time and remodeled colonic neuromuscular function by enhancing the contribution of nitrergic signaling. Mechanistically, enteric neurons emerged as direct cellular targets of f-EV-ASD, which triggered sustained Ca²⁺ elevations and increased activation of the NO pathway. Molecular profiling further revealed reduced levels of the SCFAs butyrate and valerate in f-EV-ASD compared with control EVs. These deficiencies contributed to the neuronal Ca²⁺ response through an FFAR3-dependent mechanism, identifying a potential link between EV cargo composition and enteric neuronal function. Collectively, our findings uncover a previously unrecognized f-EV-mediated pathway of microbiota-host communication that may contribute to both gastrointestinal dysfunction and gut-brain signaling abnormalities associated with ASD.

The contribution of the gut microbiota to ASD pathophysiology remains an active and evolving area of investigation^41^. Studies involving fecal microbiota transfer from individuals with ASD to mice have provided evidence that the gut microbiota can contribute to both gastrointestinal and behavioral phenotypes^16,17,42^. These effects have been proposed to involve bacterial metabolites, such as SCFAs, bile acids and phenolic compounds, several of which have been found to be altered in the stool of individuals with ASD compared with healthy controls^43,44^. Our study extends this line of research by identifying f-EVs as potential mediators of microbiota–host communication and by demonstrating that f-EVs from control individuals and patients with ASD differentially affect gut and brain functions. To date, only a limited number of studies have investigating bacterial EVs in the context of ASD. EVs derived from a single-strain probiotic *Lactobacillus paracasei* habe been shown to alleviate ASD-like behaviors in dopamine receptor D2 knockout mice^24^. Another study using metagenomic profiling of urinary EVs revealed distinct bacterial signatures in individuals with ASD compared with controls^45^, in line with our findings of differential SCFA content in f-EVs from ASD and control groups. Although we did not investigate the fate of f-EVs following oral administration, previous studies have shown that single-species bacterial EVs can cross the intestinal epithelial barriers through transcytosis^46^, enter the blood circulation, and reach various organs including the brain^47^.

Furthermore, EVs derived from pathogenic bacterial strains have been demonstrated to affect gut homeostasis, including barrier integrity and inflammation^48^, and to cross the blood brain barrier and induce neuroinflammation, neurotoxicity and cognitive impairement^49^. Together with our finding, these data highlight the importance of EV-associated gut ecosystem as an additional signaling pathway influencing gut and brain functions.

Unlike previous studies in adults with ASD without intellectual disability, which reported reduced overall bacterial diversity, we found no significant differences in alpha or beta diversity between the ASD and control groups^50,51^. This discrepancy may be partly attributable to the relatively small sample size in our study. However, at the family level, we observed a lower relative abundance of the SCFA-producing family *Lachnospiraceae*^52,53^ in the ASD group compared with the control group. Importantly, previous studies have shown that conventional analyses of gut microbiota composition may not accurately reflect its functional potential, particularly with regard to functions mediated by their EVs^27,54,55^. This highlight the importance of better characterizing the gut vesiculome to gain a more comprehensive understanding of the gut microbiota-host communication.

Consistent with previous studies^25,27,56,57^, we found that EVs isolated from human feces were from both prokaryotic and eukaryotic cellular origins as bacterial markers (OmpA, LPS, peptidoglycan, LTA, LPS) and EV markers selective for eukaryotic cell origin (CD63, CD81) were detected in the EVs derived from human feces. We found that f-EVs contained Gram negative- (OmpA and LPS) and Gram positive- (LTA) markers, indicating that both types of bacteria are able to release EVs as previously described^27,57^. The Western blot analysis performed in this study could not allow us to quantitatively determine the proportion of EVs from bacterial vs. eukaryotic origins but previous studies investigating DNA content in human fecal EVs by metagenomic analyses indicated that the vast majority of fecal EVs were from bacterial origin^27^. Even though our data support a role of the EV community within the gut ecosystem in gut and brain physiology, the respective contribution of bacterial-and host-derived EVs to these effects remains unclear and warrant further investigations following the selective purification of the bacteria-derived EVs.

A major finding of our study is that f-EVs from patients with ASD induced marked behavioral and gastrointestinal alterations in mice compared with f-EVs from control individuals. These findings indicate that f-EVs, as components of the human gut luminal microenvironment, can modulate gut and brain functions, providing new insights into microbiota–host communication mechanisms potentially involved in ASD pathophysiology.

Regarding behavior, EVs from patients with ASD induced hypersocial and anxiolytic-like behavior. The hypersocial phenotype, observed in both males and females across four independent sets of mice, was unexpected as fecal microbiota transfer from children with ASD to mice has been reported to reduce sociability compared with transfer from control children^17,58^. This apparent discrepancy may reflect differences in the functional properties of a complex bacterial community versus those of the purified f-EV fraction. At the brain protein level, f-EV-ASD reduced PSD95 expression in the cerebral cortex and cerebellum. PSD95 is a postsynaptic scaffolding protein at glutamatergic excitatory synapses that regulates the trafficking and function of glutamatergic receptors^59^. Its expression has been implicated in the regulation of social behavior as heterozygous PSD95 knockout mice display hypersociability^60,61^, potentially through NMDA-mediated excitatory synaptic current in the prefrontal cortex^62^. Thus, the hypersociability induced by f-EV-ASD may be related to reduced cerebral PSD95 expression. Interstingly, the increased sociability observed in PSD95 mutant mice has been proposed to reflect a form of ‘social blindness’, characterized by impaired processing of social cues and reduced social distance^61^. Likewise, the anxiolytic-like behavior induced by f-EV-ASD treatment might also result from an impaired ability to adequately respond to environmental cues. Although anxiety is a highly prevalent comorbidity in individuals diagnosed with ASD, previous behavioral studies in different mouse models of ASD have reported inconsistent anxiety-like phenotypes, ranging from anxiogenic to anxiolytic behavior or no significant changes^63–65^. The anxiolytic-like behavior induced by f-EV-ASD might reflect altered risk-assessment and risk-taking behaviors, as reported in the *CASPR2* KO and *SHANK3b* KO mouse models of ASD. Such alterations have been proposed to be relevant to the self-injurious behaviors reported in individuals with ASD^64^. Several limitations should nevertheless be considered. First, a broader battery of behavioral tests would help determine the specificity of the observed social and anxiety-related effects. Second, oral administration of human f-EVs to healthy mice may not fully recapitulate the chronic exposure, host genetics, and environmental factors present in ASD.

Regarding gut functions, f-EV-ASD administration markedly altered gut motility. f-EV-ASD induced an accelerated transit time and an increased EFS-induced contractile response in colonic segment, which was strongly sensitive to the NOS inhibitor L-NAME, whereas L-NAME had no effect in mice treated with f-EV-CTL. These findings suggest that f-EV-ASD affect gut motility through remodeling of the ENS nitrergic pathway. The magnitude of the nitrergic component of colonic motility positively correlated with the frequency of constipation reported by the corresponding patients, supporting a potential contribution of f-EV-ASD to dysregulated nitrergic control of colonic motility, and consequently, to constipation in ASD. Consistent with this mechanism, f-EV-ASD increased activated nNOS level, NO concentration and protein tyrosine nitration in the colon, indicative of enhanced NO signaling and nitrosative stress. These results are consistent with previous studies implicating disrupted NO signaling in ASD. In the *NL3^R451C^* mouse model of ASD, accelerated intestinal transit time was associated with an increased number of nitrergic neurons in the jejunum^10^. Conversely, pharmacological inhibition of nNOS in *Shank3^Δ4–22^* and *Cntnap2(–/–)* mouse models normalized NO production, nitrosative stress and ASD-like behavior^36,66^. In humans, increased nitrosative stress biomarkers have been reported in the blood and brain of children with ASD and in iPSC-derived cortical neurons from patients with ASD compared with controls^36,67,68^. Altogether, these findings support the potential role of NO dysregulation in ASD and raise the possibility that gut luminal EVs may act as upstream signaling mediators triggering dysregulation of NO signaling in host organs.

The mechanisms underlying f-EV-induced modulation of gut physiology remain to be fully elucidated, but our finding identified enteric neurons as direct targets of f-EVs. Indeed, f-EV-ASD increased nNOS expression and NO levels in cultured enteric neurons. The activation of nNOS is triggered by Ca^2+^ influx and Ca^2+^-calmodulin binding while its association with PSD95 promotes its anchoring to the plasma membrane near synaptic sites, facilitating rapid NO release into the perisynatic space^33,34^. Consistent with enhanced NO signaling, we found that f-EV-ASD induced sustained Ca^2+^ transients and increased the number of PSD95 synaptic clusters in nitrergic neurons compared with f-EV-CTL.

Increased PSD95 clustering may therefore favor nNOS anchoring and sustained NO release following f-EV-ASD exposure. The mechanisms implicated in the prolonged Ca^2+^ elevation induced by f-EV-ASD remain unclear but may involve differences in the content of SCFAs in f-EVs. f-EV-ASD contain lower levels of butyrate and valerate than f-EV-CTL, consistent with the lower relative abundance of *Lachnospiraceae H* in individuals with ASD, a bacterial family comprising several SCFA-producing genera, such as *Roseburia*, *Anaerostipes*, and *Coprococcus,* which are major contributors to butyrate production^53,69^. Supporting a potential role for SCFA cargo, acute supplementation with butyrate and valerate reduced the duration of f-EV-ASD-induced Ca^2+^ responses, whereas FFAR3 blockade prolonged f-EV-CTL-induced Ca^2+^ responses, reproducing the f-EV-ASD phenotype. Butyrate has well-established anti-neuroinflammatory and neuroprotective effects on enteric neurons^70–72^ and has been shown to counteract LPS-induced effects, including the anxiety and social deficits in the prenatal LPS-exposed rat model of ASD^73–75^. As LPS levels were similar in f-EV-CTL and f-EV-ASD and that LPS can activate enteric neurons ^76,77^, one possibility is that LPS-associated f-EVs contribute to Ca^2+^ neuronal response, while the lower SCFA content of f-EV-ASD reduces this inhibitory modulation, resulting in sustained Ca^2+^ elevation. Finally, the negative association between SCFA levels in f-EV-ASD and the clinical score of the corresponding patient with ASD for specific aspects of social interaction and gastrointestinal symptoms further support a potential contribution of EV-delivered SCFAs to the mechanisms underlying behavioral and gastrointestinal alterations in ASD. However, our findings do not establish that SCFAs or FFAR3 are the causal f-EV components responsible for the behavioral and nitrergic phenotypes observed *in vivo*. Other f-EV-associated molecules, such as peptidoglycans, may also contribute, either independently or in combination with SCFAs. Further studies using selective manipulation or depletion of specific f-EV cargo will therefore be required to determine the respective contribution of these components to EV-mediated effects.

One of the strengths of our translational study is the investigation of f-EVs in ASD pathophysiology within a controlled clinical setting, using individually processed samples for microbiota and metabolomics analyses, as well as for functional assays performed in mice and cultured enteric neurons. This experimental design allowed us to account for interindividual variability. However, larger, independent cohorts would be needed to validate the f-EV signature identified in this study. In conclusion, our findings supports a potential role for f-EVs as key mediators of the behavioral and GI motility alterations associated with ASD. These effects may occur through the action of f-EVs from the gut lumen on underlying intestinal tissues, including enteric neurons, as well as through more distant effects on the brain. Our data further identified the NO pathway as a key target of altered butyrate-FFAR3 signaling between f-EVs from individuals with ASD and enteric neurons.

## ACKNOWLEDGMENTS

We thank Lise Coutineau, Clémence Lalloué, Tony Durand, Thibaud Oullier, Anne Bessard, Amandine Lefebvre (UTE Animal facilities) and Steven Nedellec (platform Micropicell, SFR Francois Bonamy, Nantes). We also thank Jean-Romain Richard for the management of stool biocollection, Florence Manero from the electron microscopy platform of the University of Angers, and Sophie Brouard and Amandine Dupuy (CR2TI) for the use of the NanoSight.

## Author Contributions

BG, RA, MN, MC and HB designed the study; BG, IB, CLBS, JM, PA, ND, MM, CD, AF, LND, AT, MC performed the experiments. BG, IB, CLBS, JM, PA, CT, ND, FP, CD, AL, PE, LND, FT, JD, ST, AT, MN, MC and HB analyzed data and interpreted results; ML performed clinical evaluation and established the stool biocollection; BG, MC and HB wrote the manuscript and prepared the figures. JM, ST, AT and MN revised the manuscript. All authors read and approved the final manuscript.

## REFERENCES

1 Lai, M. C., Lombardo, M. V. & Baron-Cohen, S. Autism. Lancet 383, 896–910 (2014). 10.1016/S0140-6736(13)61539-1

2 Zeidan, J. et al. Global prevalence of autism: A systematic review update. Autism research : official journal of the International Society for Autism Research 15, 778–790 (2022). 10.1002/aur.2696

3 Coury, D. L. et al. Gastrointestinal Conditions in Children With Autism Spectrum Disorder: Developing a Research Agenda. Pediatrics 130, S160–S168 (2012). 10.1542/peds.2012-0900N

4 Ferguson, B. J., Dovgan, K., Takahashi, N. & Beversdorf, D. Q. The Relationship Among Gastrointestinal Symptoms, Problem Behaviors, and Internalizing Symptoms in Children and Adolescents With Autism Spectrum Disorder. Frontiers in Psychiatry 10, 194 (2019). 10.3389/fpsyt.2019.00194

5 Furness, J. B. Types of neurons in the enteric nervous system. J Auton Nerv Syst 81, 87–96 (2000). 10.1016/s0165-1838(00)00127-2

6 Hotta, R. et al. Transplanted ENSCs form functional connections with intestinal smooth muscle and restore colonic motility in nNOS-deficient mice. Stem Cell Research & Therapy 14, 232 (2023). 10.1186/s13287-023-03469-3

7 Johnson, C. D. et al. Deletion of *choline acetyltransferase* in enteric neurons results in postnatal intestinal dysmotility and dysbiosis. The FASEB Journal 32, 4744–4752 (2018). 10.1096/fj.201701474RR

8 Sasselli, V. et al. Planar cell polarity genes control the connectivity of enteric neurons. Journal of Clinical Investigation 123, 1763–1772 (2013). 10.1172/JCI66759

9 Golubeva, A. V. et al. Microbiota-related Changes in Bile Acid & Tryptophan Metabolism are Associated with Gastrointestinal Dysfunction in a Mouse Model of Autism. EBioMedicine 24, 166–178 (2017). 10.1016/j.ebiom.2017.09.020

10 Hosie, S. et al. Gastrointestinal dysfunction in patients and mice expressing the autism-associated R451C mutation in neuroligin-3. Autism Research 12, 1043–1056 (2019). 10.1002/aur.2127

11 Gonzales, J. et al. Fecal Supernatant from Adult with Autism Spectrum Disorder Alters Digestive Functions, Intestinal Epithelial Barrier, and Enteric Nervous System. Microorganisms 9, 1723 (2021). 10.3390/microorganisms9081723

12 Hughes, H. K., Rose, D. & Ashwood, P. The Gut Microbiota and Dysbiosis in Autism Spectrum Disorders. Current Neurology and Neuroscience Reports 18, 81 (2018). 10.1007/s11910-018-0887-6

13 Liu, F. et al. Altered composition and function of intestinal microbiota in autism spectrum disorders: a systematic review. Translational Psychiatry 9, 43 (2019). 10.1038/s41398-019-0389-6

14 Peralta-Marzal, L. N. et al. A robust microbiome signature for autism spectrum disorder across different studies using machine learning. Scientific Reports 14, 814 (2024). 10.1038/s41598-023-50601-7

15 Xu, M., Xu, X., Li, J. & Li, F. Association Between Gut Microbiota and Autism Spectrum Disorder: A Systematic Review and Meta-Analysis. Frontiers in Psychiatry 10, 473 (2019). 10.3389/fpsyt.2019.00473

16 Prince, N. et al. Mouse strain-specific responses along the gut-brain axis upon fecal microbiota transplantation from children with autism. Gut Microbes 17, 2447822 (2025). 10.1080/19490976.2024.2447822

17 Xiao, L. et al. Fecal Microbiome Transplantation from Children with Autism Spectrum Disorder Modulates Tryptophan and Serotonergic Synapse Metabolism and Induces Altered Behaviors in Germ-Free Mice. mSystems 6, e01343–01320 (2021). 10.1128/mSystems.01343-20

18 Van Niel, G. et al. Challenges and directions in studying cell–cell communication by extracellular vesicles. Nature Reviews Molecular Cell Biology 23, 369–382 (2022). 10.1038/s41580-022-00460-3

19 Villard, A., Boursier, J. & Andriantsitohaina, R. Bacterial and eukaryotic extracellular vesicles and nonalcoholic fatty liver disease: new players in the gut-liver axis? Am J Physiol Gastrointest Liver Physiol 320, G485–G495 (2021). 10.1152/ajpgi.00362.2020

20 Villard, A., Boursier, J. & Andriantsitohaina, R. Microbiota-derived extracellular vesicles and metabolic syndrome. Acta Physiol (Oxf) 231, e13600 (2021). 10.1111/apha.13600

21 Choi, J., Kim, Y.-K. & Han, P.-L. Extracellular Vesicles Derived from *Lactobacillus plantarum* Increase BDNF Expression in Cultured Hippocampal Neurons and Produce Antidepressant-like Effects in Mice. Experimental Neurobiology 28, 158–171 (2019). 10.5607/en.2019.28.2.158

22 Haas-Neill, S. & Forsythe, P. A Budding Relationship: Bacterial Extracellular Vesicles in the Microbiota-Gut-Brain Axis. Int J Mol Sci 21 (2020). 10.3390/ijms21238899

23 Kang, C.-s., et al. Extracellular Vesicles Derived from Gut Microbiota, Especially Akkermansia muciniphila, Protect the Progression of Dextran Sulfate Sodium-Induced Colitis. PLoS ONE 8, e76520 (2013). 10.1371/journal.pone.0076520

24 Park, J.-Y. et al. Lactobacillus paracasei-derived extracellular vesicles reverse molecular and behavioral deficits in mouse models of autism spectrum disorder. Experimental & Molecular Medicine 57, 788–803 (2025). 10.1038/s12276-025-01429-w

25 Fizanne, L. et al. Faeces-derived extracellular vesicles participate in the onset of barrier dysfunction leading to liver diseases. Journal of Extracellular Vesicles 12, 12303 (2023). 10.1002/jev2.12303

26 Ruiz-Malagón, A. J. et al. An Exploratory Study on the Pathogenic Role of Faecal Extracellular Vesicles in Metabolic Dysfunction-Associated Steatotic Liver Disease Progression and in Drug-Induced Liver Injury. Journal of Extracellular Vesicles 15, e70240 (2026). 10.1002/jev2.70240

27 Zhu, L. T. et al. Diverse functional genes harboured in extracellular vesicles from environmental and human microbiota. Journal of Extracellular Vesicles 11, 12292 (2022). 10.1002/jev2.12292

28 Ghozlane, A. et al. Accurate profiling of microbial communities for shotgun metagenomic sequencing with Meteor2. Microbiome 13, 227 (2025). 10.1186/s40168-025-02249-w

29. PMAC, S. M. o. Large Targeted Metabolomics - LC-MS Parameters of Q-Exactive - MetaboHUB-Tours V.2 (2026). 10.17504/protocols.io.eq2lyoq9qgx9/v2

30 Monteiro, J., Lefèvre, A, Dufour-Rainfray, D, Oury, A, Chicheri, G, Galineau, L, Blasco, H, Nadal-Desbarats, L, and Emond, P. Multi-Compartment SCFA Quantification in Humans. American Journal of Analytical Chemistry 15, 177–200 (2024). 10.4236/ajac.2024.156012

31 Caillaud, M. et al. A functional network of highly pure enteric neurons in a dish. Frontiers in Neuroscience 16, 1062253 (2023). 10.3389/fnins.2022.1062253

32 Donovan, A. P. A. & Basson, M. A. The neuroanatomy of autism – a developmental perspective. Journal of Anatomy 230, 4–15 (2017). 10.1111/joa.12542

33 Brenman, J. E. et al. Interaction of nitric oxide synthase with the postsynaptic density protein PSD-95 and alpha1-syntrophin mediated by PDZ domains. Cell 84, 757–767 (1996). 10.1016/s0092-8674(00)81053-3

34 Chaudhury, A., He, X.-D. & Goyal, R. K. Role of PSD95 in membrane association and catalytic activity of nNOSα in nitrergic varicosities in mice gut. American Journal of Physiology-Gastrointestinal and Liver Physiology 297, G806–G813 (2009). 10.1152/ajpgi.00279.2009

35 Rameau, G. A., Chiu, L.-Y. & Ziff, E. B. Bidirectional regulation of neuronal nitric-oxide synthase phosphorylation at serine 847 by the N-methyl-D-aspartate receptor. The Journal of Biological Chemistry 279, 14307–14314 (2004). 10.1074/jbc.M311103200

36 Tripathi, M. K. et al. The NO Answer for Autism Spectrum Disorder. Advanced Science 10, 2205783 (2023). 10.1002/advs.202205783

37 Kuo, W. N., Kreahling, J. M., Shanbhag, V. P., Shanbhag, P. P. & Mewar, M. Protein nitration. Molecular and Cellular Biochemistry 214, 121–129 (2000). 10.1023/a:1007118300731

38 Chachlaki, K. & Prevot, V. Nitric oxide signalling in the brain and its control of bodily functions. British Journal of Pharmacology 177, 5437–5458 (2020). 10.1111/bph.14800

39 Nohr, M. K. et al. GPR41/FFAR3 and GPR43/FFAR2 as cosensors for short-chain fatty acids in enteroendocrine cells vs FFAR3 in enteric neurons and FFAR2 in enteric leukocytes. Endocrinology 154, 3552–3564 (2013). 10.1210/en.2013-1142

40 Tsao, P. & Von Zastrow, M. Downregulation of G protein-coupled receptors. Current Opinion in Neurobiology 10, 365–369 (2000). 10.1016/S0959-4388(00)00096-9

41 Qin, S. et al. Research progress on the microbiota-gut-brain axis in autism spectrum disorder: a narrative review. Front Microbiol 17, 1824546 (2026). 10.3389/fmicb.2026.1824546

42 Sharon, G. et al. Human Gut Microbiota from Autism Spectrum Disorder Promote Behavioral Symptoms in Mice. Cell 177, 1600–1618 e1617 (2019). 10.1016/j.cell.2019.05.004

43 Bu, W., Chen, Z., Liu, B. & Jia, X. Gut microbiota and its metabolism in autism spectrum disorder: from pathogenesis to therapy. Front Cell Infect Microbiol 15, 1687691 (2025). 10.3389/fcimb.2025.1687691

44 Ganachaud, B. et al. Interaction of the Gut Microbiota and the Enteric Nervous System: Implications in Gastrointestinal Disorders Associated With Autism Spectrum Disorder. Autism Res 19, e70328 (2026). 10.1002/aur.70328

45 Lee, Y. et al. Rapid Assessment of Microbiota Changes in Individuals with Autism Spectrum Disorder Using Bacteria-derived Membrane Vesicles in Urine. Experimental neurobiology 26, 307–317 (2017). 10.5607/en.2017.26.5.307

46 Rubio, A. P. D. et al. Transcytosis of Bacillus subtilis extracellular vesicles through an in vitro intestinal epithelial cell model. Scientific reports 10, 3120 (2020). 10.1038/s41598-020-60077-4

47 Kang, M., Jordan, V., Blenkiron, C. & Chamley, L. W. Biodistribution of extracellular vesicles following administration into animals: A systematic review. J Extracell Vesicles 10, e12085 (2021). 10.1002/jev2.12085

48 Yi, H., Li, M., Xu, L. & Mao, H. Deciphering the immunomodulatory cross-talk: Bacterial extracellular vesicles in gut homeostasis. Virulence 16, 2566255 (2025). 10.1080/21505594.2025.2566255

49 Kaisanlahti, A. et al. Bacterial extracellular vesicles - brain invaders? A systematic review. Front Mol Neurosci 16, 1227655 (2023). 10.3389/fnmol.2023.1227655

50 Gonzales, J. et al. Fecal Supernatant from Adult with Autism Spectrum Disorder Alters Digestive Functions, Intestinal Epithelial Barrier, and Enteric Nervous System. Microorganisms 9 (2021). 10.3390/microorganisms9081723

51 Ying, J. et al. The Gut Microbiota in Young Adults with High-Functioning Autism Spectrum Disorder and Its Performance as Diagnostic Biomarkers. Nutrients 17 (2025). 10.3390/nu17111748

52 Abdugheni, R. et al. Metabolite profiling of human-originated Lachnospiraceae at the strain level. Imeta 1, e58 (2022). 10.1002/imt2.58

53 Louis, P. & Flint, H. J. Formation of propionate and butyrate by the human colonic microbiota. Environ Microbiol 19, 29–41 (2017). 10.1111/1462-2920.13589

54 Kaisanlahti, A. et al. Gut microbiota-derived extracellular vesicles form a distinct entity from gut microbiota. mSystems 10, e00311–00325 (2025). 10.1128/msystems.00311-25

55 Rodríguez-Díaz, C. et al. The Metagenomic Composition and Effects of Fecal-Microbe-Derived Extracellular Vesicles on Intestinal Permeability Depend on the Patient’s Disease. International Journal of Molecular Sciences 24, 4971 (2023). 10.3390/ijms24054971

56. Byts, N. et al. in Cell-Secreted Vesicles Vol. 2668 (ed Seppo Vainio) 211–226 (Springer US, 2023).

57 Kameli, N., Borman, R., López-Iglesias, C., Savelkoul, P. & Stassen, F. R. M. Characterization of Feces-Derived Bacterial Membrane Vesicles and the Impact of Their Origin on the Inflammatory Response. Frontiers in Cellular and Infection Microbiology 11, 667987 (2021). 10.3389/fcimb.2021.667987

58 Avolio, E. et al. Modifications of Behavior and Inflammation in Mice Following Transplant with Fecal Microbiota from Children with Autism. Neuroscience 498, 174–189 (2022). 10.1016/j.neuroscience.2022.06.038

59 Kim, E. & Sheng, M. PDZ domain proteins of synapses. Nature Reviews Neuroscience 5, 771–781 (2004). 10.1038/nrn1517

60 Feyder, M. et al. Association of Mouse *Dlg4* (PSD-95) Gene Deletion and Human *DLG4* Gene Variation With Phenotypes Relevant to Autism Spectrum Disorders and Williams’ Syndrome. American Journal of Psychiatry 167, 1508–1517 (2010). 10.1176/appi.ajp.2010.10040484

61 Winkler, D. et al. Hypersocial behavior and biological redundancy in mice with reduced expression of PSD95 or PSD93. Behavioural Brain Research 352, 35–45 (2018). 10.1016/j.bbr.2017.02.011

62 Coley, A. A. & Gao, W.-J. PSD-95 deficiency disrupts PFC-associated function and behavior during neurodevelopment. Scientific Reports 9, 9486 (2019). 10.1038/s41598-019-45971-w

63 Dhamne, S. C. et al. Replicable in vivo physiological and behavioral phenotypes of the Shank3B null mutant mouse model of autism. Molecular Autism 8, 26 (2017). 10.1186/s13229-017-0142-z

64 Oron, O. et al. Gene network analysis reveals a role for striatal glutamatergic receptors in dysregulated risk-assessment behavior of autism mouse models. Translational Psychiatry 9, 257 (2019). 10.1038/s41398-019-0584-5

65 Yang, M. et al. Reduced excitatory neurotransmission and mild autism-relevant phenotypes in adolescent Shank3 null mutant mice. The Journal of Neuroscience: The Official Journal of the Society for Neuroscience 32, 6525–6541 (2012). 10.1523/JNEUROSCI.6107-11.2012

66 Bazbaz, W., Kartawy, M., Khaliulin, I. & Amal, H. Nitric oxide inhibition ameliorates cortical proteomic changes in the Cntnap2-/- and Shank3Δ4-22 mouse models of autism spectrum disorder. Molecular Autism 17, 21 (2026). 10.1186/s13229-026-00716-1

67 Dong, D., Zielke, H. R., Yeh, D. & Yang, P. Cellular stress and apoptosis contribute to the pathogenesis of autism spectrum disorder. Autism Research 11, 1076–1090 (2018). 10.1002/aur.1966

68 Söğüt, S. et al. Changes in nitric oxide levels and antioxidant enzyme activities may have a role in the pathophysiological mechanisms involved in autism. Clinica Chimica Acta 331, 111–117 (2003). 10.1016/S0009-8981(03)00119-0

69 Fitzgerald, B. G. et al. Members of Lachnospiraceae produce valerate and caproate in response to short-chain fatty acids. Microbiome 13, 201 (2025). 10.1186/s40168-025-02220-9

70 Brossaud, R. et al. The short-chain fatty acid butyrate prevents gut-brain amyloid-β pathology and neuroinflammation in an Alzheimer mouse model. Molecular Psychiatry (2026). 10.1038/s41380-026-03522-6

71 Caetano, M. A. F., Magalhães, H. I. R., Duarte, J. R. L., Conceição, L. B. & Castelucci, P. Butyrate Protects Myenteric Neurons Loss in Mice Following Experimental Ulcerative Colitis. Cells 12, 1672 (2023). 10.3390/cells12131672

72 Soret, R. et al. Short-Chain Fatty Acids Regulate the Enteric Neurons and Control Gastrointestinal Motility in Rats. Gastroenterology 138, 1772–1782.e1774 (2010). 10.1053/j.gastro.2010.01.053

73 Huuskonen, J., Suuronen, T., Nuutinen, T., Kyrylenko, S. & Salminen, A. Regulation of microglial inflammatory response by sodium butyrate and short-chain fatty acids. British Journal of Pharmacology 141, 874–880 (2004). 10.1038/sj.bjp.0705682

74 Qiu, J. et al. Lipopolysaccharide-Induced Depression-Like Behaviors Is Ameliorated by Sodium Butyrate via Inhibiting Neuroinflammation and Oxido-Nitrosative Stress. Pharmacology 105, 550–560 (2020). 10.1159/000505132

75 Wang, X. et al. Sodium butyrate facilitates CRHR2 expression to alleviate HPA axis hyperactivity in autism-like rats induced by prenatal lipopolysaccharides through histone deacetylase inhibition. mSystems, e00415–00423 (2023). 10.1128/msystems.00415-23

76 Breunig, E. et al. Histamine excites neurones in the human submucous plexus through activation of H1, H2, H3 and H4 receptors. The Journal of Physiology 583, 731–742 (2007). 10.1113/jphysiol.2007.139352

77 Coquenlorge, S. et al. Modulation of lipopolysaccharide-induced neuronal response by activation of the enteric nervous system. Journal of Neuroinflammation 11, 202 (2014). 10.1186/s12974-014-0202-7

