## Supplemental figures for "Fecal extracellular vesicles from adults with autism spectrum disorder affect mouse behavior and gut motility by targeting the neuronal nitrergic pathway"

### SUPPLEMENTARY DATA

| Metabolite class | Name of metabolites | CTL | ASD | p-value |
| --- | --- | --- | --- | --- |
| Amino Acids-Peptides-Derivatives | 4-HYDROXY-L-PHENYLGLYCINE | 0.00142119 | 0.00239502 | 0.50885 |
|  | ALPHA-AMINOADIPATE | 0.00154408 | 0.00155833 | 0.66297 |
|  | BETAINE | 0.03341462 | 0.050068 | 0.42002 |
|  | CARNOSINE | 0.00299223 | 0.00224323 | 0.64108 |
|  | CITRULLINE | 0.00566946 | 0.005384 | 0.33619 |
|  | CREATINE | 0.00452123 | 0.00573063 | 0.27855 |
|  | GLYCINE | 0.00171923 | 0.00148453 | 0.64104 |
|  | L-ALANINE | 0.02166154 | 0.01786 | 0.13048 |
|  | L-ARGININE | 0.11766154 | 0.094549 | 0.37098 |
|  | L-ASPARAGINE | 0.01286077 | 0.0100403 | 0.55814 |
|  | L-ASPARTATE | 0.00680769 | 0.00573097 | 0.30747 |
|  | LEUCINE | 0.04517692 | 0.03454 | 0.56215 |
|  | L-GLUTAMIC ACID | 0.0361 | 0.03387 | 0.10402 |
|  | L-GLUTAMINE | 0.148 | 0.12100667 | 0.78164 |
|  | L-HISTIDINE | 0.00961692 | 0.0084436 | 0.1068 |
|  | LL-2,6-DIAMINOHEPTANEDIOATE | 0.01688462 | 0.03604379 | 0.6961 |
|  | L-LYSINE | 0.01117692 | 0.00874967 | 0.12738 |
|  | L-METHIONINE | 0.07013077 | 0.06016786 | 0.92972 |
|  | L-PHENYLALANINE | 0.01016538 | 0.0090494 | 0.8501 |
|  | L-PIPECOLIC ACID | 0.00328454 | 0.00260435 | 0.71483 |
|  | L-PROLINE | 0.02695231 | 0.02727167 | 0.36433 |
|  | L-SERINE | 0.01526923 | 0.01321367 | 0.62293 |
|  | L-THREONINE | 0.00621692 | 0.004769 | 0.55374 |
|  | L-TRYPTOPHAN | 0.01878077 | 0.01602357 | 0.44679 |
|  | L-TYROSINE | 0.00459231 | 0.00394465 | 0.85193 |
|  | L-VALINE | 0.02368385 | 0.02410375 | 0.16961 |
|  | N6,N6,N6-Trimethyl-L-lysine | 0.11390769 | 0.15395 | 0.22166 |
|  | N-ACETYL-DL-GLUTAMIC ACID | 0.00114123 | 0.00186773 | 0.10407 |
|  | N-ACETYL-L-LEUCINE | 0.00142631 | 0.00221039 | 0.17171 |
| Organic acids | 2-Methylbutyrylglycine | 0.00075938 | 0.0012169 | 0.079818 |
|  | **3-Methyladipic acid** | **0.01124846** | **0.01748233** | **0.030223\*** |
|  | AZELAIC ACID | 0.34738462 | 0.4362 | 0.18171 |
|  | **Succinic acid** | **0.00890015** | **0.03777403** | **0.004769\*** |
|  | Urocanic acid | 0.00538654 | 0.00430573 | 0.95981 |
| Phosphoric acids | DIHYDROXYACETONE PHOSPHATE | 0.09312308 | 0.08477 | 0.95039 |
|  | GLYCERALDEHYDE 3-PHOSPHATE<br>DIETHYL ACETAL | 0.01166308 | 0.036921 | 0.288 |
| Acyl-carnitines | L-Carnitine | 0.00641615 | 0.00729633 | 0.23446 |
|  | Octadecanoylcarnitine | 0.00293892 | 0.0035902 | 0.87149 |
| Cyclic and acyclic aliphatic compounds | ACETYLCHOLINE | 0.00070582 | 0.0009619 | 0.56221 |
|  | CADAVERINE | 0.0198575 | 0.02765167 | 0.22166 |
|  | CREATININE | 0.00369192 | 0.00490948 | 0.69611 |
|  | CYCLOPENTANONE | 0.00200023 | 0.00284587 | 0.36209 |
| Aromatic compounds-Vitamins-Amines | (S)-1-PHENYLETHANOL | 0.01049846 | 0.01340433 | 0.76235 |
|  | ADENINE | 0.03081154 | 0.0353718 | 0.8501 |
|  | BENZOATE | 0.04710133 | 0.03805621 | 0.93974 |
|  | BILIVERDIN | 0.00109197 | 0.0017434 | 0.56911 |
|  | CYTOSINE | 0.00552592 | 0.00382328 | 0.49294 |
|  | GUANINE | 0.02256231 | 0.01764833 | 0.1582 |
|  | HYPOXANTHINE | 0.13726154 | 0.1024 | 0.30746 |
|  | LUMICHROME | 0.00847269 | 0.00558489 | 0.8501 |
|  | NICOTINATE | 0.005528 | 0.0075995 | 0.57505 |
|  | TRANS-CINNAMALDEHYDE | 0.01636846 | 0.02556367 | 0.79132 |
|  | TRIGONELLINE | 0.002701 | 0.00490957 | 0.40565 |
|  | TYRAMINE | 0.00495385 | 0.00709233 | 0.13379 |
|  | XANTHINE | 0.00312843 | 0.00292196 | 0.7143 |
|  | **12-hydroxy-9-cis-octadecenoic acid** | **0.10997692** | **0.02890267** | **0.0062508\*** |
| Lipids: Fatty acids. Steroids. Hormones. Vitamins | Brassicasterol | 0.01174615 | 0.01522533 | 0.91971 |
|  | DEOXYCARNITINE | 0.00799231 | 0.02064766 | 0.49294 |
|  | Deoxycholic acid | 0.12375342 | 0.0848734 | 0.73371 |
|  | Dodec-2-enedioic acid | 0.01219538 | 0.00888 | 0.075651 |
|  | Lithocholic acid | 0.0303 | 0.02386626 | 0.18586 |
|  | LPC(16_0) | 0.00259723 | 0.01682513 | 0.37772 |
|  | LPC(18_0) | 0.00189008 | 0.00785487 | 0.40565 |
|  | LPE(16_0) | 0.00074251 | 0.00103277 | 0.77198 |
|  | METHYL JASMONATE | 0.00809846 | 0.00828867 | 0.35762 |
|  | PC(32_0) | 0.00043672 | 0.0011026 | 0.87982 |
|  | PC(34_1) | 0.00085152 | 0.00165251 | 0.38466 |
|  | PC(34_2) | 0.00061141 | 0.0005388 | 0.61427 |
|  | SM(34_0) | 0.00067069 | 0.00056484 | 0.41281 |
|  | SM(34_1) | 0.00144431 | 0.00295633 | 1 |
|  | SN-GLYCERO-3-PHOSPHOCHOLINE | 0.00192485 | 0.00325527 | 0.13378 |
| Nucleotides. Nucleosides. and Derivatives | 5'-DEOXYADENOSINE | 0.00127575 | 0.00302011 | 0.35765 |
|  | ADENOSINE | 0.01152938 | 0.01509167 | 0.288 |
|  | CYTIDINE | 0.00512531 | 0.00385183 | 0.23446 |
|  | **INOSINE** | **0.00386231** | **0.0024729** | **0.021129\*** |
| Sugars | N-ACETYL-D-GLUCOSAMINE | 0.01148746 | 0.02294533 | 0.22166 |
|  | N-ACETYLNEURAMINATE | 0.04652246 | 0.07104533 | 0.41276 |

Table S1: Metabolomics analysis reveals differential concentration of metabolite between f-EV-ASD and f-CTL-EV, n=13 for CTL and n= 27 for ASD. Analysis was performed using the MetaboAnalyst 6.0 platform. Data are mean ± SEM. p-values are calculated using a t-test. n=13 CTL, n=28 ASD.

|  | CTL | ASD | p-value |
| --- | --- | --- | --- |
| **TRYPTOPHAN** | 24203±7994 n=13 | 14353±2365 n=24 | 0,09 |
| **KYNURENINE** | 5.235±0.4945 n=13 | 4.54±0.60<br>n=24 | 0,1 |
| **5-OH-TRYPTOPHANE** | 32.1±16.7 n=4 | 16.46±8.22 n=5 | 0,85 |
| **INDOLE-3-ALDEHYDE** | 7.81±1.40 n=13 | 6.92±1.49 n=21 | 0,22 |
| **INDOLE-3-ACETIC ACID** | 5.25±2.13 n=14 | 3.48±0.74 n=12 | 0,78 |

Table S2: Table showing the concentrations of Tryptophan derivatives present in f-EV-ASD and f-EV-CTL. Data are represented as the mean ± SEM in nanomolar (nM). All p-values are calculated using the Mann-Whitney test according to the normality test.

**A**

**Three-chamber test**

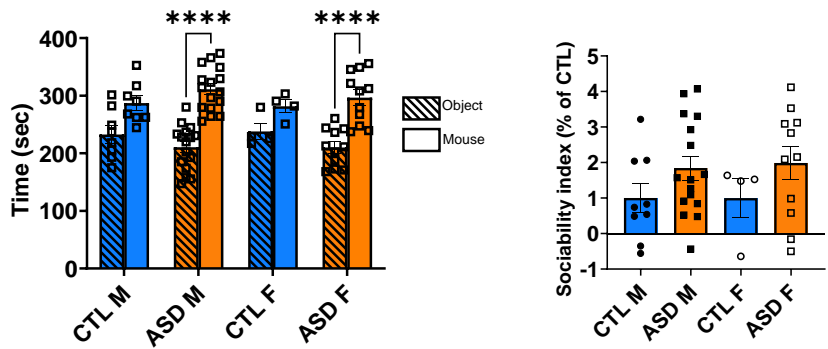

**B**

**Elevated plus maze**

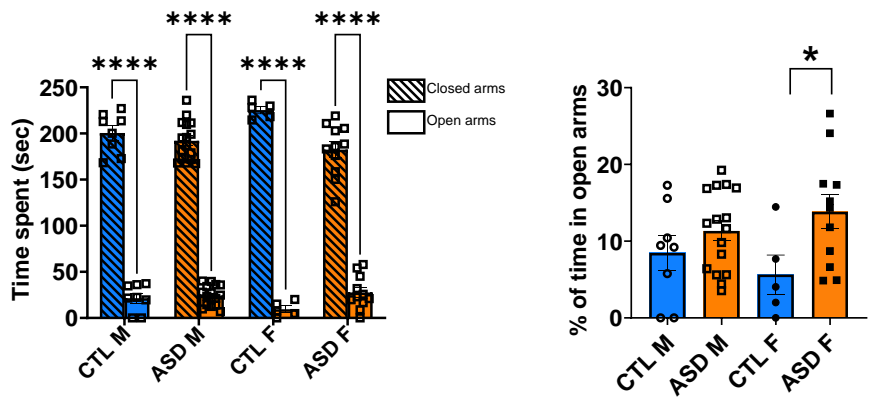

**C**

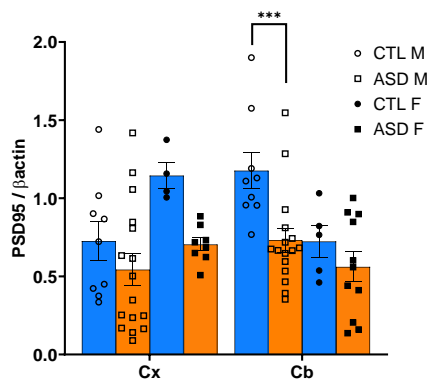

**Supplemental Figure S1: Effects of f-EV treatment according to the sex of the donor and of the recipient mouse.** (A,B) Behavioral assessments in males (M) and females (F) using the three-chamber test (A) and the elevated plus maze (B). (C) PSD95 expression in the cortex and hippocampus of male and female mice treated with f-EV-CTL or f-EV-ASD from male and female donor respectively. Data are presented as mean ± SEM with individual values shown. \*$p < 0.05$, \*\*\*$p < 0.001$, \*\*\*\*$p < 0.0001$, Two-way ANOVA test, $n = 8$ CTL males, $n = 16$ ASD males, $n = 4-5$ CTL females, $n = 11$ ASD females.

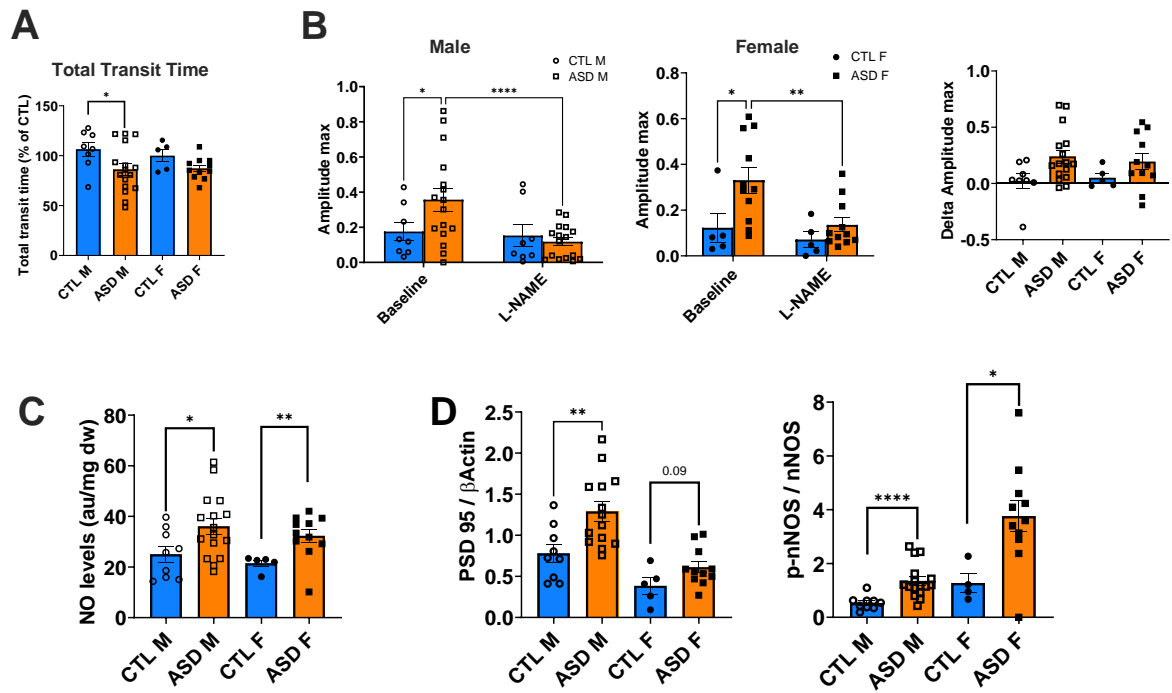

**Supplemental Figure S2: Effects of f-EV treatment according to the sex of the donor and of the recipient mouse in GI functions.** (A) *In vivo* total transit time expressed as percentage of the control group (CTL) in male (M) and female (F) mice. (B) *Ex vivo* colonic motility in male and female mice treated with f-EV-CTL or f-EV-ASD from male and female donors respectively. Maximum amplitude of the post-EFS response of colon segments from f-EV-CTL- and f-EV-ASD-treated mice in basal condition and after sequential addition of L-Name (50 μM) and atropine (1 μM). \*\*\*\*$p$<0.0001, \*\*$p$<0.01, \*$p$<0.05, 2-way ANOVA repeated measure with Bonferroni post hoc test, $n$=8 CTL males, $n$=16 ASD males, $n$=5 CTL females, $n$=11 ASD females. Right panel: the L-Name-sensitive maximum amplitude (difference between maximum amplitude in basal condition and in the presence of L-Name) in colon segments from f-EV-CTL- and f-EV-ASD-treated male and female mice. $p$>0.05, Mann-Whitney test. (C) NO concentration in the colon of f-EV-CTL- and f-EV-ASD-treated male and female mice. (D) Protein expression levels normalized to β-actin signal in the colon of mice treated with f-EV-CTL or f-EV-ASD for PSD95 and p-nNOS at Ser1412. For C and D: \*$p$<0.05, \*\*$p$<0.01, Mann-Whitney test, $n$=9 CTL males, $n$=16 ASD males, $n$=5 CTL females, $n$=11 ASD females.

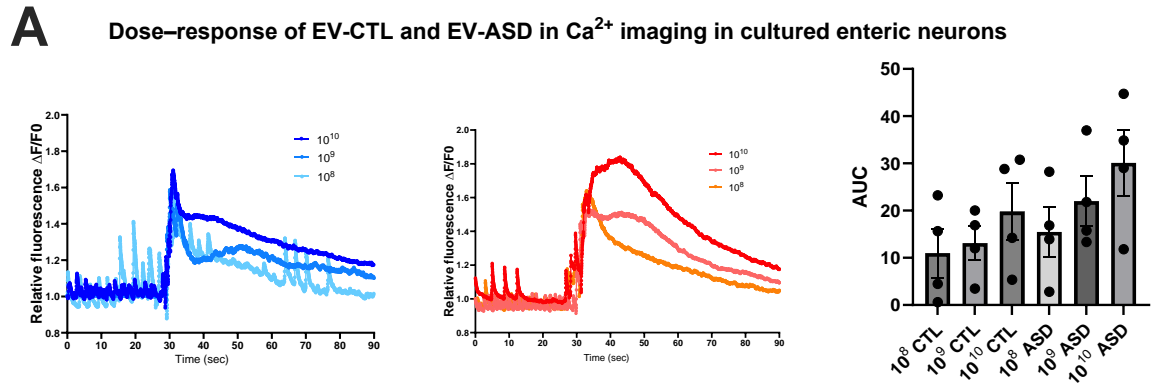

**B** Spontaneous $\text{Ca}^{2+}$ transients in enteric neurons after 48h treatment with EVs

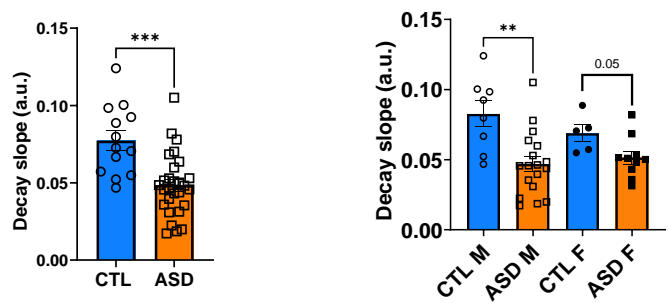

**C** Evoked intracellular $\text{Ca}^{2+}$ elevation in enteric neurons upon EV stimulation

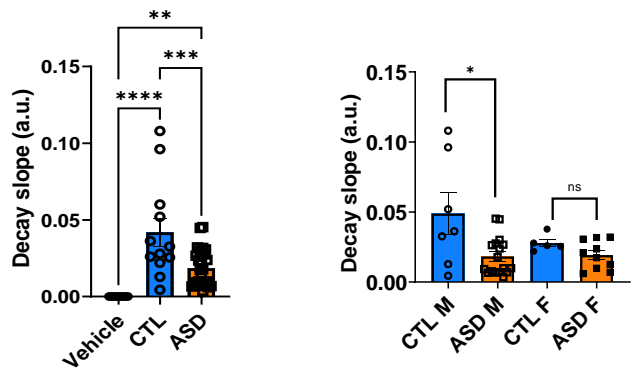

**Supplemental Figure S3: Effects of f-EV-ASD and f-EV-CTL on evoked and spontaneous $\text{Ca}^{2+}$ transients in enteric neurons.** (A) Dose-response of f-EV-CTL and f-EV-ASD from 3 distinct individuals on $\text{Ca}^{2+}$ transients in cultured enteric neurons. Right panel: quantification of the area under the curve (AUC) of the evoked $\text{Ca}^{2+}$ peak. (B) Quantification of the decay slope of spontaneous $\text{Ca}^{2+}$ transients in enteric neurons treated for 48h with f-EV-CTL f- EV-ASD from male (M) or female (F) individuals. \*\*$p < 0.01$, \*\*\*$p = 0.001$, Welch's t-test. (C) Quantification of evoked intracellular $\text{Ca}^{2+}$ responses for the decay slopes in cultured enteric neurons upon stimulation with PBS, f-EV-CTL or f-EV-ASD from male or female individuals. \*$p < 0.05$ and \*\*$p < 0.01$, \*\*\*$p < 0.001$, \*\*\*\*$p < 0.0001$, ANOVA and Mann-Whitney test.
